# A potent and selective TNKS2 inhibitor for tumor-selective WNT suppression

**DOI:** 10.1101/2025.03.04.641305

**Authors:** Jill Zimmerman, Brandon F Malone, Efrat Finkin-Groner, Shan Sun, Rui Liang, Miguel Foronda, Emma M Schatoff, Elizabeth Granowsky, Sukanya Goswami, Alyna Katti, Benjamin Leach, Heather Alcorn, Tuomas Tammela, Yoshiyuki Fukase, Tanweer Khan, David J Huggins, John Ginn, Nigel Liverton, Richard K Hite, Lukas E Dow

## Abstract

Hyperactive WNT signaling is a potent cancer driver, but clinical translation of WNT inhibitors has been hampered by on-target toxicities. WNT signaling can be constrained through inhibition of the PARP family enzymes Tankyrase 1 (TNKS1) and Tankyrase 2 (TNKS2), however, existing TNKS inhibitors suppress WNT signaling in both tumor and healthy tissues. In this study, we show that the loss of chromosome 8p that occurs in approximately half of advanced epithelial malignancies, creates a collateral vulnerability that enables tumor-selective inhibition of Tankyrase activity. 8p loss depletes expression of TNKS1 and creates a tumor-specific dependency on the functionally redundant TNKS2 protein. Through structure-guided drug design, we identify a first-in-class TNKS2-selective inhibitor that can drive selective WNT inhibition in TNKS1-deficient oncogenic cell and organoid models. This work demonstrates a targetable vulnerability in multiple cancer types, providing a new approach to potent and selective WNT-targeted therapies.

## INTRODUCTION

Hyperactivation of the WNT pathway is a near-ubiquitous feature of colorectal cancer (CRC). Suppressing WNT hyperactivation through genetic or pharmacologic approaches can prevent tumor growth and/or drive sustained tumor regression (1-8). WNT activation is also implicated in the pathogenesis of numerous other tumor types, including breast, lung, prostate, gastric, ovarian and hepatocellular cancers (9), drives therapy resistance (10-13), and can suppress activity of immune checkpoint inhibitors (14, 15). Thus, while there is substantial clinical potential for targeting WNT signaling in cancer, early clinical studies, and in vivo pre-clinical work with WNT inhibitors have revealed significant on-target toxicities in normal tissues (16, 17).

One strategy to suppress WNT signaling is through inhibition of Tankyrase activity. Tankyrase (TNKS; hereafter TNKS1 for clarity) and Tankyrase 2 (TNKS2) are functionally redundant PARP-family enzymes that promote WNT signaling through the PARylation and degradation of the negative regulator AXIN1 (18) and have been shown to modulate YAP signaling by targeting AMOT proteins (19). Many small molecule compounds have been developed that target TNKS1/TNKS2 enzymes and these compounds can effectively stabilize AXIN1, downregulate WNT signaling, and suppress tumor cell proliferation in multiple cancer types (16-18, 20-23). However, like other broad WNT inhibitors, potent TNKS1/ TNKS2 inhibitors cause dose-limiting tissue toxicity in vivo, limiting their therapeutic potential (16, 17).

During malignant progression, many cancers acquire large-scale chromosomal amplifications and deletions. One of the most frequent genomic alterations in human epithelial cancers are deletions on the short arm of chromosome 8 (8p), which contains the TNKS1 locus (24-28). Here, we show that tumors with 8p deletions, which express reduced or no TNKS1 protein, carry an acquired tumor-specific dependence on TNKS2 for cellular Tankyrase activity. In these cells, WNT pathway activity is regulated primarily by TNKS2 and can be diminished by selective suppression of TNKS2. Through rational drug design, we develop a first-in-class TNKS2-selective small molecule inhibitor that suppresses the growth of TNKS1-deleted but not TNKS1-diploid cells. This work represents the first description of a targetable collateral vulnerability in the WNT pathway and demonstrates the feasibility for the development of potent and selective TNKS2 inhibitors as a therapeutic modality.

## RESULTS

### Chromosome 8p deletions sensitize cells to TNKS2-selective silencing

Large genomic deletions in Chromosome 8 are frequently observed in many epithelial cancer types. Analysis of available ICGC and TCGA whole genome sequencing data (29) shows approximately half of colon, lung, breast, liver, and prostate cancers carry heterozygous or homozygous 8p deletions (Fig. 1A). Loss of this genomic segment and reduced TNKS1 gene dosage correlates with a decrease in TNKS1 transcript and protein expression (Fig. 1B-C, fig. S1A-B). Consistent with previously published data (18, 30), suppression of both TNKS1 and TNKS2 in 8p diploid DLD1 cells was required to stabilize AXIN1; however consistent with an induced dependency on TNKS2 in 8p-deleted cells, silencing of TNKS2 alone in HCC1171 cells was sufficient to stabilize AXIN1 (Fig. 1D).

**Figure 1:**
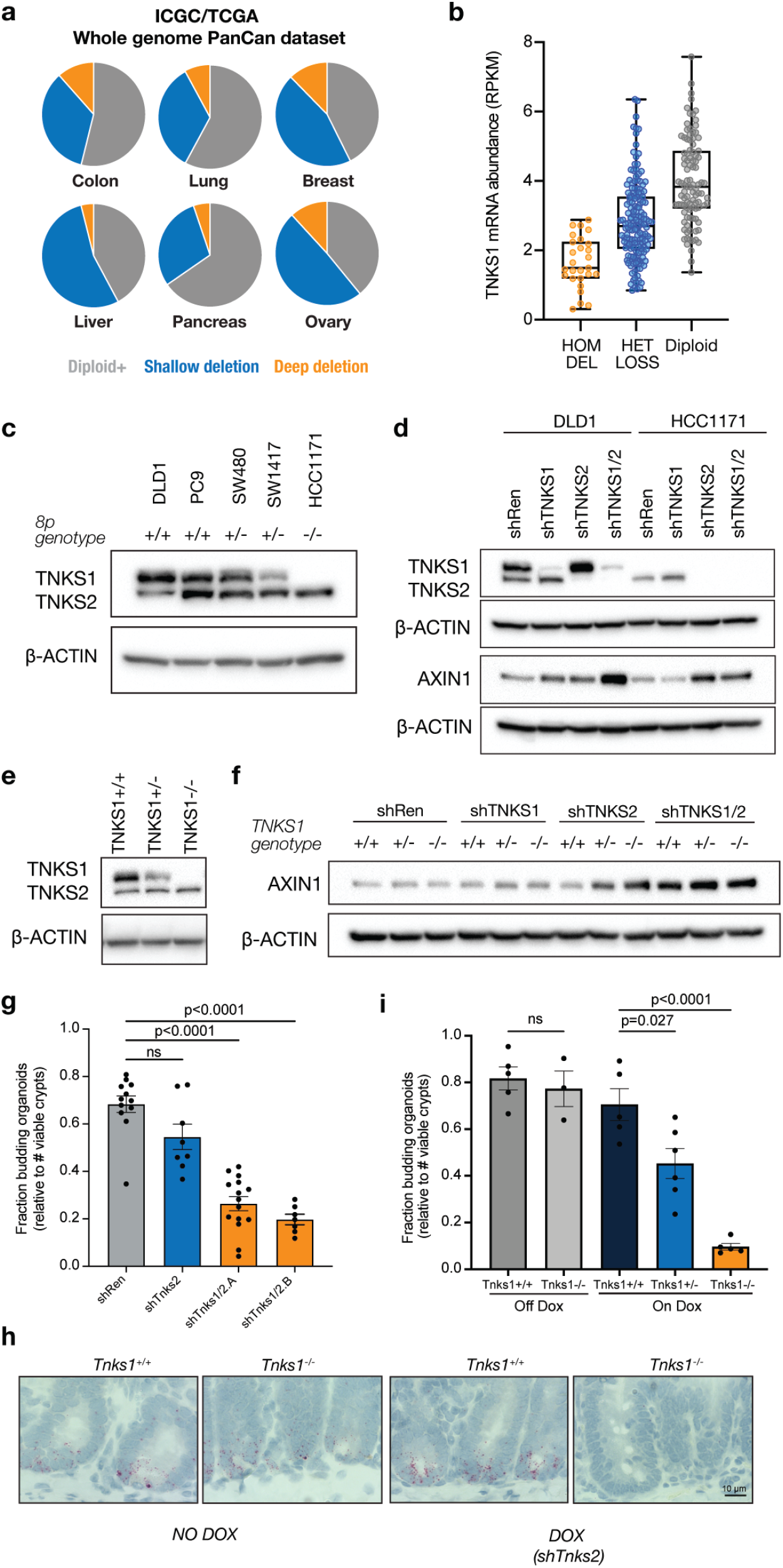
8p deletion and TNKS1 loss sensitizes cells to TNKS2-selective suppression. (A) Fraction of common epithelial cancers carrying heterozygous or homozygous 8p deletions. Data is derived from the ICGC/TGCA PanCan dataset. (B) TNKS1 genomic copy number is correlated with TNKS1 transcript expression in CRC. Data is derived from the ICGC/TCGA PanCan dataset. Center line, median; box limits, upper and lower quartiles; whiskers, min to max. (C) Western blot showing reduced TNKS1 protein expression with 8p deletions in a panel of colorectal (DLD1, SW480, SW1417) and lung (PC9, HCC1171) cancer cell lines. (D) Western blot showing AXIN1 and TNKS1/2 levels following TNKS1, TNKS2, or TNKS1/2 suppression. shTNKS2 induced AXIN1 stabilization in 8p null cells (HCC1171) but not 8p diploid cells (DLD1). (E) Western blot showing TNKS1 protein expression in TNKS1WT, TNKS1Het, and TNKS1KO DLD1s. (F) Western blot showing AXIN1 levels following TNKS1, TNKS2, or TNKS1/2 suppression. shTNKS2 induced AXIN1 stabilization in DLD1 TNKS1Het and TNKS1KO isogenic clones but not TNKS1WT cells. (G) Fraction of viable organoids 4 days after crypt isolation from mice expressing shRNAs for two weeks. shTnks1/2.A and shTnks1/2.B represent two different shRNA combinations (n = 7-15, mean with SEM, one-way ANOVA with Tukey correction). (H) In situ hybridization of Lgr5 expression in the small intestine of Tnks1WT or Tnks1KO mice on dox (shTnks2) or off dox (no shRNA). (I) Fraction of viable organoids 4 days after crypt isolation from mice on dox (shTnks2) or off dox (no shRNA) for three weeks. (n = 3-6, mean with SEM, one-way ANOVA with Tukey correction).

The 8p chromosome arm contains several potential tumor suppressor genes whose aggregate loss may promote tumorgenicity (25, 28). To confirm the selective effect of TNKS2 suppression is specifically due to the loss of TNKS1, we generated homozygous knockout (TNKS1^KO^) and heterozygous (TNKS1^HET^) isogenic DLD1 clones (Fig. 1E). As expected, in TNKS1^KO^ cells, TNKS2 silencing induced AXIN1 stabilization to a level equivalent to suppression of both TNKS enzymes (Fig. 1F). Importantly, AXIN1 stabilization was also apparent at intermediate levels in TNKS1^HET^ cells, suggesting that TNKS1 is haploinsufficient for supporting cellular Tankyrase activity and that heterozygous TNKS1 loss can sensitize cells to TNKS2 inhibition (Fig. 1F).

Homozygous deletion of both Tankyrase enzymes in mice causes embryonic lethality, while TNKS2 knockout mice are viable and fertile (31). This suggests that systemic, selective TNKS2 targeting would have minimal toxicity to normal tissues. To directly test this idea in vivo, we generated mice harboring an inducible shRNA targeting Tnks2 and compared it to our previously published Tnks1/Tnks2 dual knockdown mice (3). To explore the effect of Tnks2-selective suppression on tissue function with high WNT pathway activity in vivo, we measured the ability of intestinal crypts to form budding organoids in Matrigel culture. Silencing of both Tnks1 and Tnks2 induced a significant decrease in stem-cell driven organoid formation compared to neutral (shRen) controls (Fig. 1G, p<0.01), while Tnks2 suppression showed a mild but not significant decrease in organoid formation (Fig. 1G; p>0.05).

To determine whether loss of one or both copies of Tnks1 would create a Tnks2 dependence in vivo, we used Cas9 and tandem sgRNAs to delete the entire Tnks1 locus in fertilized zygotes and intercrossed the resulting animals to generate wildtype (Tnks1^WT^), heterozygous (Tnks1^HET^) and homozygous (Tnks1^KO^) knockout mice carrying the inducible shTnks2 cassette (Fig. S2A-C, Fig. S3A). In the absence of doxycycline (no Tnks2 silencing) there was no difference in organoid forming capacity or the expression of the WNT target Lgr5 between Tnks1^WT^ and Tnks1^KO^ crypts (Fig. 1H-I), implying Tnks1 is largely dispensable for crypt function. Similarly, Tnks1^WT^/shTnks2 mice treated with dox for 3 weeks showed robust Lgr5 expression and organoid formation comparable to untreated controls (Fig. 1H-I, Fig. S3B). In contrast, dox-treated Tnks1^KO^/shTnks2 mice showed a significant defect in organoid formation and had a marked reduction in Lgr5 expression in crypt stem cells (Fig. 1H-I). Further, consistent with the dose dependent AXIN1 stabilization seen in isogenic DLD1 cells, Tnks1^HET^/shTnks2 animals showed reduced organoid forming capacity and intermediate Lgr5 expression in intestinal crypts (Fig. 1I, Fig. S3B).

Together, these data support the concept that the loss of either one or both copies of TNKS1 following 8p deletions, creates a TNKS2 dependence that could be exploited to enable tumor cell-selective suppression of WNT signaling in 8p-deleted tumor cells while avoiding on-target toxicities associated with WNT blockade in normal tissues.

### Rational design of TNKS2-selective small molecule inhibitors

More than 50 small molecule Tankyrase inhibitors have been described in the literature (16-18, 20-23, 32). These compounds bind either the adenosine (e.g. G007-LK) (16) or nicotinamide (e.g. XAV939) (18) subsites of the enzyme (Fig. 2B), or act as dual-binders, contacting both subsites (e.g. TNKS656) (22). Most TNKS inhibitors potently inhibit both TNKS1 and TNKS2 enzymes, though some have been reported to have 20-50 fold selectivity for TNKS2 (33-35). To directly test whether these existing compounds could provide potent and selective TNKS2 inhibition we resynthesized and tested four reported TNKS2-selective molecules using a commercial PARylation assay for TNKS1 and TNKS2 (Fig. S4). In contrast to the published data, each of the previously described compounds showed less than 10-fold selectivity toward TNKS2, like reported pan-TNKS inhibitors (Fig. S4).

**Figure 2:**
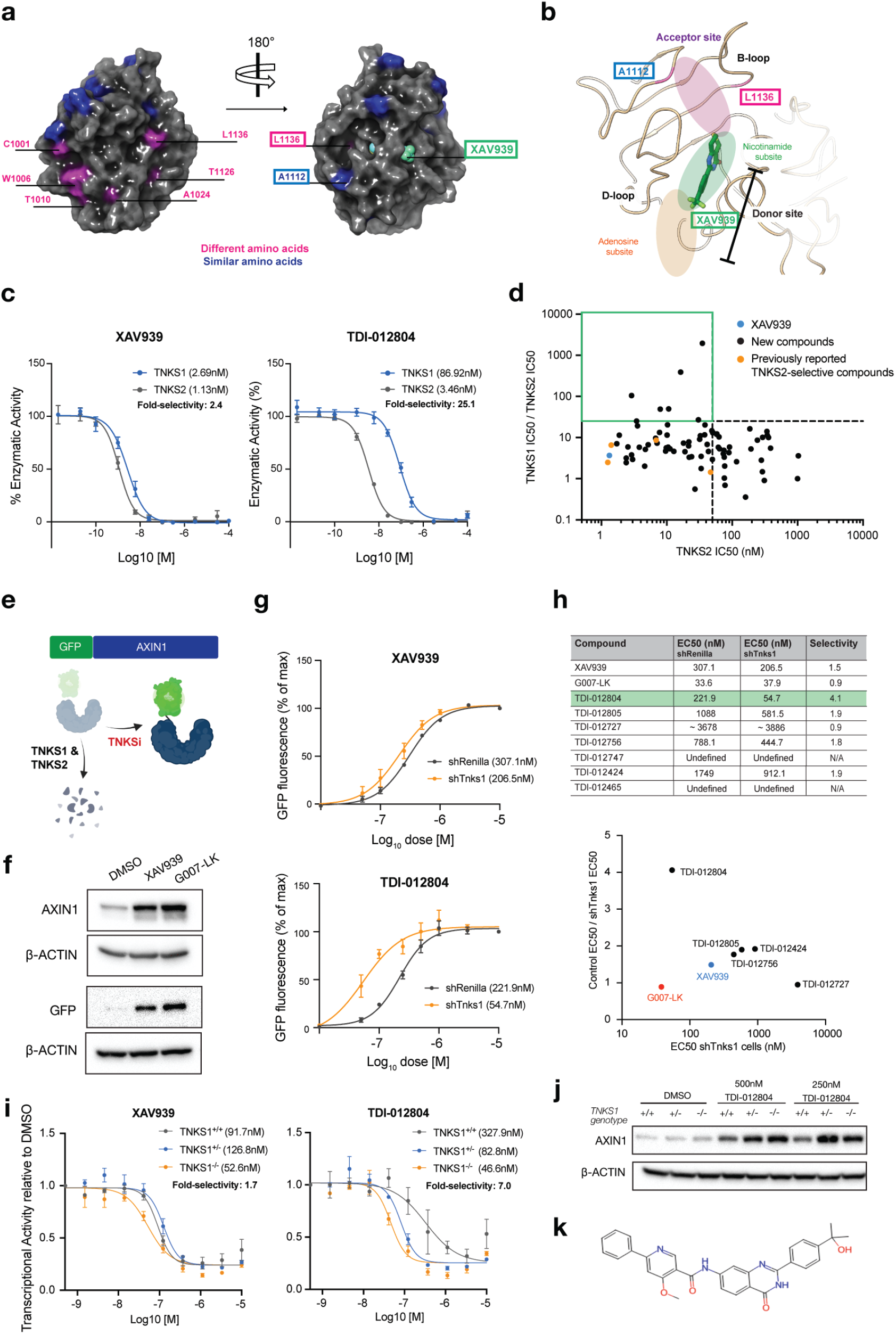
Development of TNKS2-selective small molecule inhibitors. (A) Surface displacement of TNKS2 catalytic domain with non-conserved residues highlighted. (B) XAV939 bound to the TNKS2 donor site. (C) Dose response curves of in vitro PARylation assay measuring the enzymatic activity of TNKS1 and TNKS2 incubated with XAV939 or TDI-012804 (n = 4, error bars = SEM). Enzymatic activity is normalized to DMSO. (D) Scatterplot showing the TNKS2 IC50 and TNKS2 selectivity (TNKS1 IC50 / TNKS2 IC50) of synthesized compounds (N = 2-4). Green box highlights potent (TNKS2 IC50 < 50nM) and selective (>25-fold) compounds. Blue point = XAV939. Orange points = previously published TNKS2-selective compounds. Black points = novel compounds. (E) AXIN1-GFP reporter construct to measure stabilization of AXIN1 protein. (F) Western blot of 3T3 AXIN1-GFP reporter cells showing stabilization of AXIN1 and GFP protein following incubation with 1uM XAV939 or 1uM G007-LK for 24h. (G) AXIN1-GFP stabilization following incubation with XAV939 or TDI-012804 for 24h (n = 3, error bars = SEM). GFP stabilization is quantified as the increase in fluorescence over DMSO and normalized to the highest concentration. (H) EC50 values in shRenilla and shTnks1 AXIN1-GFP reporter cells of the seven hit compounds, XAV939, and G007-LK (N=3). Selectivity is shRenilla EC50 / shTnks1 EC50. (I) Dose response curves of transcriptional activity measured using a TOPFlash reporter expressed in DLD1s (n = 3 [XAV939] or 2 [TDI-012804], error bars = SEM). Luciferase is normalized to DMSO. (J) Western blot showing selective AXIN1 stabilization in DLD1s with TDI-012804 treatment (500nM or 250nM for 24h). (K) Structure of TDI-012804. Image in (E) created with Biorender.com

The catalytic PARP domains of TNKS1 and TNKS2 have high sequence identity, particularly in the residues that surround the nicotinamide and adenosine subsites (Fig. 2A), making selective targeting of these regions challenging. Through analysis of available crystal structures of the TNKS1 and TNKS2 enzymatic domains, we noted two residues (A1112 and L1136 in TNKS2) surrounding the acceptor site that were not conserved between TNKS1 and TNKS2 (Fig. 2A). We reasoned that extending a ligand from the adjacent nicotinamide subsite into the acceptor site, could impart selectivity toward TNKS2 (Fig. 2B). We first established an initial set of molecules derived from XAV939 with quinazolin-4(3H)-one scaffold that effectively inhibited TNKS1/TNKS2 activity and then used structure-based design with docking and validated free energy calculations with FEP+ (Fig. S5A-B) to extend them toward the variant residues through the addition of substituents to quinazolin-4(3H)-one scaffold (Fig. 2B).

To identify TNKS2 selective inhibitors we undertook an SAR campaign exploring different trajectories on the quinazolin-4(3H)-one core with the objective of interacting with the flexible B-loop. In total, we synthesized over 300 small molecules with different chemical moieties and measured inhibitory activity of each using biochemical PARylation and cellular assays. From initial screening, we identified 88 small molecules with good potential potency and/or selectivity for full dose response assays against TNKS1 and TNKS2. Of the 88 small molecules in secondary screening, we identified 7 with high potency against TNKS2 (IC50 < 50nM) and greater than 25-fold selectivity toward TNKS2 (TNKS1 IC50 / TNKS2 IC50 > 25) (Fig. 2C-D, fig. S6, Table S1).

To determine whether the seven identified compounds showed selective inhibition of endogenous TNKS2 protein in cells, we quantified the stability of a GFP-AXIN1 reporter in both wildtype (TNKS1^WT^) and TNKS1 knockdown (TNKS1^KD^), NIH3T3 cells (Fig. 2E-F, fig. S7A-B); This reporter allows direct and simple quantification of AXIN1 protein stabilization while avoiding confounding changes in AXIN1 transcription. In this assay, TDI-012804 was most potent and showed the clearest selectivity profile (Fig. 2G-H, Fig. S7C). To further evaluate TDI-012804 activity in cells, we quantified the impact on downstream WNT-mediated transcription in TNKS1 isogenic DLD1 cells using an integrated TOP-Flash reporter. Consistent with the GFP-AXIN1 data, TDI-012804 was potent (EC50 <100nM) and showed TNKS2-selectivity in these cells as compared to the nonselective XAV939 (Fig. 2I, Fig. S8). As observed in TNKS2 shRNA experiments (Fig. 1F), TNKS1^HET^ cells showed increased response to TDI-012804 compared to TNKS1^WT^ cells (Fig. 2I), further supporting the notion that a single allele of TNKS1 is not sufficient to support cellular Tankyrase function.

Consistent with the GFP-AXIN1 reporter, TDI-012804 induced marked stabilization of endogenous AXIN1 in both TNKS1^HET^ and TNKS1^KO^ cells but induced only a minimal increase in AXIN1 protein in TNKS1 diploid controls (Fig. 2J). Importantly, TDI-012804 (Fig. 2K) showed no inhibition of related PARP-family enzymes (PARP1, PARP2, and PARP3), highlighting it as a potent and selective TNKS2 inhibitor (Fig. S9).

### Structural basis for TNKS2 selective inhibition

To define the mechanism underlying TNKS2 inhibition by TDI-012804 we sought to structurally characterize the ligand bound state to the TNKS2 catalytic domain. As previously described (36, 37), the minimal active ADP-ribosyl transferase unit of TNKS2, the SAM-PARP domains, adopts a helical filamentous architecture, aiding the determination of a high-resolution structure (Fig. 3A). Using a combination of helical reconstruction and focused refinements, we determined a ∼2.2 Å nominal resolution cryo-electron microscopy (cryo-EM) map of TDI-012804 bound to TNKS2 (Fig. S10, S13, Table S2). This structure illustrated that the ligand occupied both the nicotinamide donor site and the acceptor site (Fig. 3B), consistent with the computational design. TDI-012804 has 2-phenyl-3,4-dihydroquinazoline-4-one core, which is anchored in the nicotinamide subsites by similar interactions as previously reported for this scaffold. The lactam portion of the quinazolinone ring forms hydrogen bond networks with Ser1068 side chain and the Gly1032 backbone amide. The quinazolinone core forms a ∏-stacking interaction with Tyr1071, while the phenyl group at 2-position fits into a hydrophobic pocket. In the acceptor subsite, the amide NH forms a hydrogen bond with Glu1138, while the pyridine nitrogen interacts with Met1054 through water-mediated interaction. (Fig. 3C).

**Figure 3:**
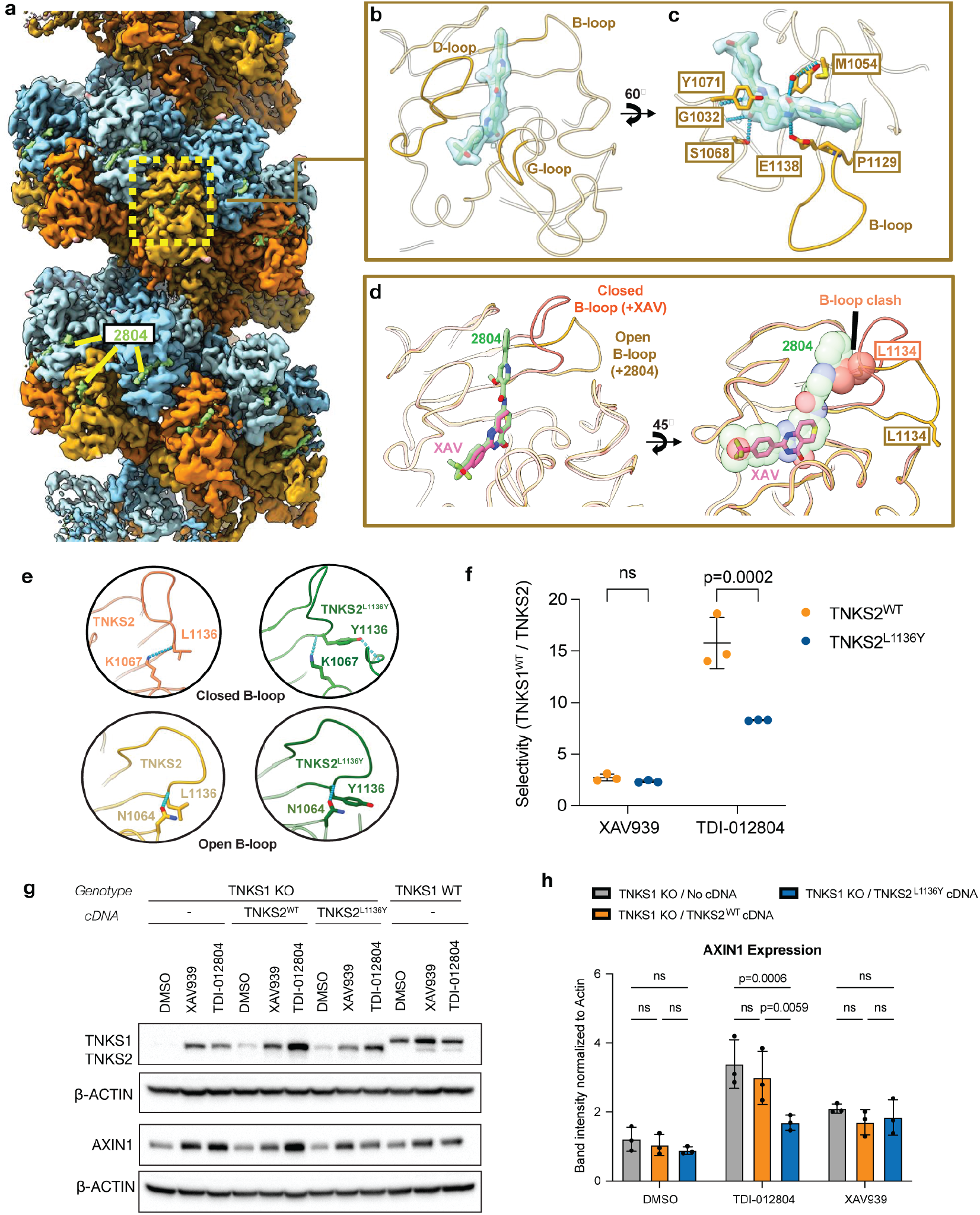
Mechanism of TNKS2-selective small molecule inhibitors. (A) Volume overview, shown as an isosurface, depicting the TNKS2 SAM-PARP filament and associated TDI-012804 binding sites (colored lime green). (B) The TDI-012804 binding site in relation to conserved tankyrase active-site motifs. TDI-012804 is shown as sticks and overlaid with the cryo-EM map density (C) Active-site residues, involved in the recognition of TDI-012804, are depicted as sticks with associated hydrogen bonds shown as dashed cyan connections. (D) Superimposition of the +XAV (ligand shown in sticks and colored pink) sand +2804 (ligand shown in sticks and colored green) models. (E) Cartoon schematics of the B-loop as observed in the experimental structures; +XAV/apo (top-left), +2804 (bottom left) and models depicting the L1136Y mutant in the closed (top right - modeled using PDB 4W6E) and open conformations. Intramolecular hydrogen bonds are depicted as cyan dashes. (F) Fold selectivity of IC50 values of XAV939 and TDI-012804 comparing the effect on TNKS1^WT^ to TNKS2^WT^ and TNKS2^L1136Y^ enzymatic activity using an in vitro PARylation assay (n = 3, mean with SD, two-way ANOVA with Sidak correction). (G) Western blot showing AXIN1 and TNKS1/2 levels in TNKS^KO^ DLD1s expressing no cDNA, TNKS2^WT^ cDNA, or TNKS2L1136Y cDNA, compared to TNKS1^WT^ DLD1s. Cells were treated with DMSO, 250nM XAV939, or 250nM TDI-012804 for 24h. (H) Quantification of AXIN1 western blot band intensity normalized to Actin (n = 3, mean with SD, two-way ANOVA with Tukey correction).

To gain insight into potential mechanisms of TNKS2 selectivity, we additionally resolved cryo-EM structures of TNKS2 in the absence of ligand (apo) and bound to the non-selective inhibitor XAV939 (Fig. S11-13). Analysis of these structures revealed that the terminal phenyl ring of TDI-012804 forces the B-loop in the acceptor subsite to adopt an “open” conformation in contrast to the “closed” conformation observed in the presence of XAV939 or without ligand (Fig. 3D). While the TNKS2 B-loop is notably dynamic in all conditions, the closed loop conformation of Leucine 1134 resolved in the presence of XAV939 or in the absence of ligand would clash with the phenyl ring in TDI-012804, reducing ligand association (Fig. S14A-D). However, rearrangement of the B-loop into the open conformation when bound to TDI-012804 re-positions L1134 away from the active site and allows the phenyl ring to form a pi-pi stacking interaction with P1129. An analysis of all known crystal structures of both TNKS1 and TNKS2 indicates that the TNKS1 B-loop is modeled in the closed conformation near exclusively whereas the TNKS2 B-loop has been modeled in a continuum of conformations between the open and closed conformations.

Together, these data indicate that the B-loop is more dynamic in TNKS2 than in TNKS1. A single amino acid (L1136 of TNKS2; Y1289 in TNKS1) distinguishes the B-loops of TNKS1 and TNKS2; We reasoned that the enhanced intramolecular hydrogen bonding network provided by a tyrosine in this position may increase the rigidity of the B-loop of TNKS1 and thus impede binding of TDI-012804 (Fig. 3E). To directly test this, we purified TNKS2^WT^, TNKS2^L1136Y^ and TNKS1 SAM-PARP domains and measured the IC50 for TDI-012804 and XAV939 by *in vitro* PARylation assay. Consistent with our model, mutation of L1136 to tyrosine (L1136Y) significantly reduced the potency of TDI-012804 against TNKS2 but had no effect on the activity of XAV939 (Fig. 3F, Fig. S15A). These data suggest that L1136 in TNKS2 contributes to the enhanced flexibility of the TNKS2 B-loop, thereby increasing TDI-012804 affinity. To investigate the effect of the L1136Y substitution on the selectivity of TDI-012804 in cells, we generated TNKS1^KO^ DLD1 cell lines that exogenously expressed TNKS2^WT^ or TNKS2^L1136Y^ and measured AXIN1 stabilization 24h following treatment. Consistent with the biochemical assay, TDI-012804 increased AXIN1 protein in TNKS1^KO^ and TNKS1^KO^/TNKS2^WT^ cells, but not in TNKS1^WT^ or TNKS1^KO^/TNKS2^L1136Y^ cells (Fig. 3G-H, Fig. S15B). Consistent with the PARylation assay, the L1136Y substitution did not alter the response to the pan-TNKS inhibitor XAV939 (Fig. 3G-H, Fig. S15B).

### TNKS2-selective inhibitors suppress WNT signaling in TNKS-depleted cells

Tankyrase inhibition in DLD1 cells induces WNT pathway suppression but does not impact cell proliferation or viability (16, 30). To test the effect of TDI-012804 in a WNT-dependent system that closely mimics normal intestinal biology, we generated Tnks1^WT^, Tnks1^HET^ and Tnks1^KO^ intestinal organoid cultures harboring a homozygous nonsense mutation (Q1405X) in the APC tumor suppressor (Fig 4A-B). This mutation models common truncating events within the mutation cluster region of APC observed in both familial and sporadic CRC. Consistent with previously published strong WNT and TNKS dependence of APC mutant organoids (3), RNA-sequencing of G007-LK treated Apc^Q1405X^/Tnks1^WT^ and Apc^Q1405X^/Tnks1^KO^ small intestinal organoids revealed broad transcriptional changes including reduced stem and progenitor cell signatures and upregulation of intestinal differentiation markers across multiple cell lineages (Fig. 4C-E). Apc^Q1405X^/ Tnks1^KO^ organoids treated with TDI-012804 showed a near-identical overall transcriptional response to G007-LK treated cultures, while Apc^Q1405X^/Tnks1^WT^ organoids showed no significant gene expression changes compared to DMSO controls (Fig. 4C-E). Notably, there were only two significantly deregulated genes in the RNAseq analysis comparing G007-LK and TDI-012804 in Tnks1^KO^ organoids (Table S3), implying that TDI-012804 does not have measurable non-Tankyrase (off-target) activity at effective doses.

**Figure 4:**
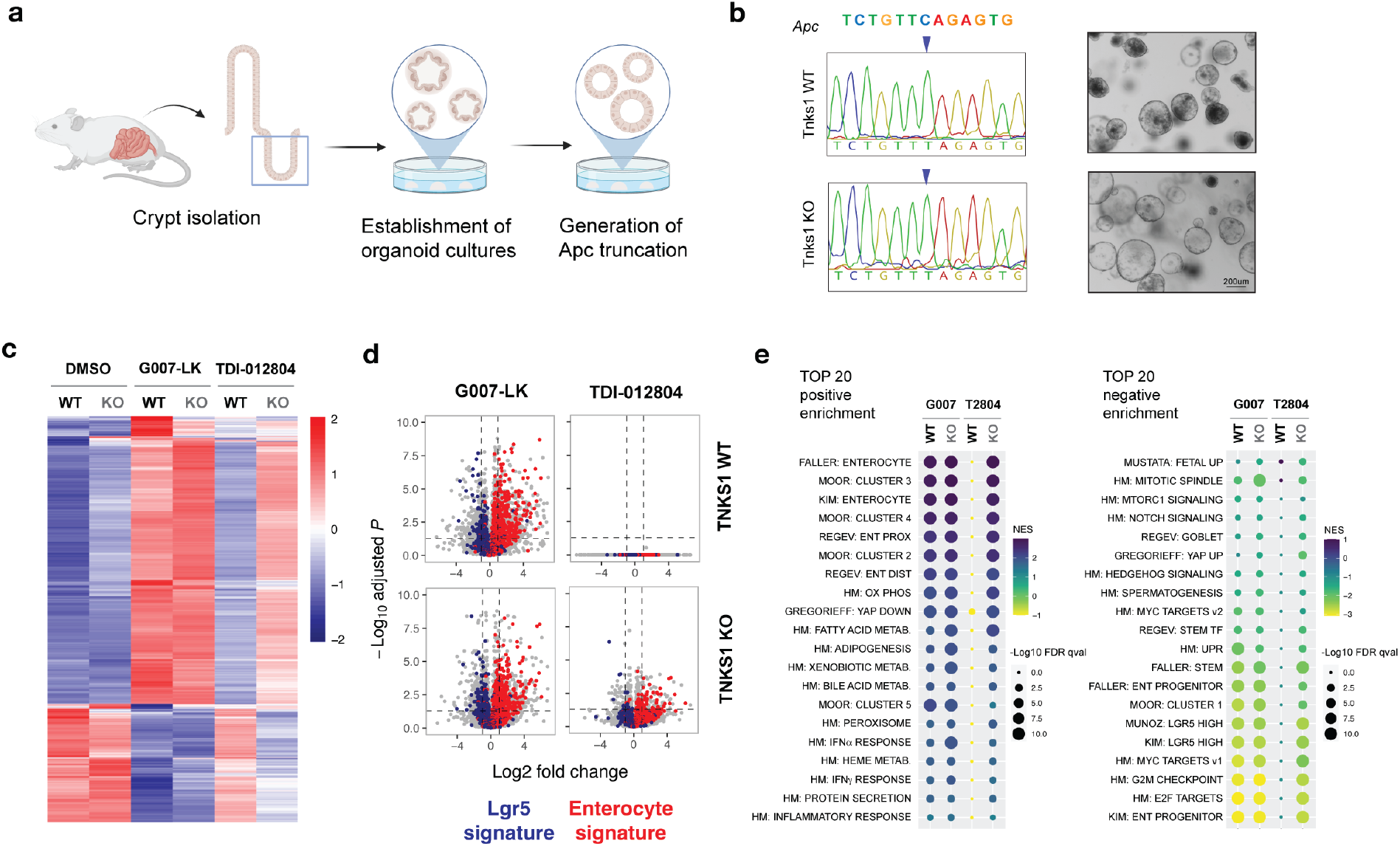
TNKS2-selective inhibitors suppress WNT in TNKS-depleted cells. (A) Generation of small intestinal organoid lines carrying ApcQ1405X mutations. (B) C>T mutations in Apc allele in organoid lines, and brightfield images of Apc mutant organoids. Scale bar = 200um. (C) Heatmap showing average log2 normalized expression of all differentially expressed genes (log2 fold change > 1, padj<0.05) between G007-LK and DMSO treatments (n=3 replicates/condition). (D) Volcano plots highlighting average log2 fold change of genes enriched in Lgr5-high cells (“Lgr5 signature”) or differentiated enterocytes (“Enterocyte signature”), comparing treatments as indicated to DMSO treated organoids (n=3/ condition). (E) Summary of the top 20 positively and negatively enriched gene sets enriched (from the G007-LK vs DMSO comparison) ranked by normalized enrichment score (NES). Plot shows NES (color) and false discovery rate qval (size) for all treatment conditions compared to DMSO treated cells of the same genotype. Image in (A) created with Biorender.com

### TNKS2-selective inhibitors block proliferation in TNKS-depleted cancer cells and organoids

To further explore the effects of TNKS2 inhibition on cell survival and proliferation, we identified several human cancer cell lines that are responsive to TNKS inhibition either alone or in combination with other targeted therapies (16, 30, 38). These included one 8p heterozygous cell line, SW403 (CRC), and two 8p diploid lines, OVCAR4 (Ovarian) and H23 (lung). Consistent with the response seen in TNKS1 heterozygous DLD1 cells, TDI-012804 treatment of SW403 cells induced AXIN1 stabilization to similar levels as the pan-TNKS inhibitor G007-LK (Fig. 5A). Consequently, treatment with either TDI-012804 or G007-LK resulted in a similar reduction in colony formation (Fig. 5B). As expected, treatment of 8p diploid H23 cells with TDI-012804 did not induce AXIN1 stabilization nor suppress colony formation like G007-LK (Fig. S16A); however, suppression of TNKS1 in these cells induced a significant sensitization to TDI-012804 that was further enhanced in the presence of Palbociclib (Fig. 5B, Fig. S16B). OVCAR4 cells were sensitive to pan-TNKS inhibition by G007-LK with or without Palbociclib (Fig. 5B, Fig. S16A-B), and consistent with the response seen in H23 cells, TDI-012804 significantly reduced OVCAR4 colony formation only in TNKS1-depleted cells (Fig. 5B).

**Figure 5:**
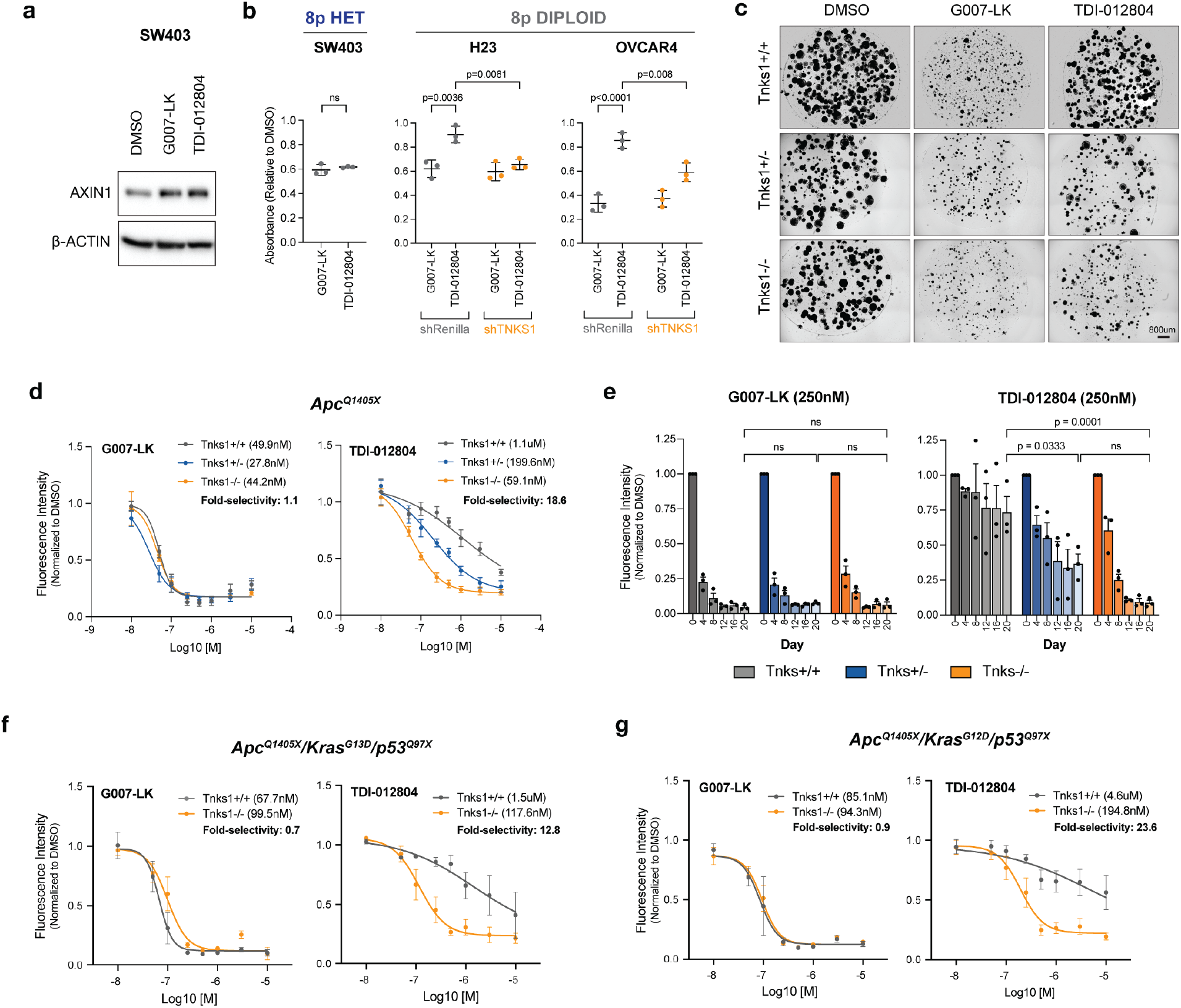
TNKS2 inhibitors reduce proliferation and viability in TNKS1-depleted cells. (A) Western blot showing stabilization of AXIN1 protein levels in SW403 cells treated with G007-LK (250nM) or TDI-012804 (250nM) for 24h. (B) Quantification of SW403 colony forming assays treated with 250nM G007-LK or 250nM TDI-012804 normalized to DMSO (n = 3, mean with SD, unpaired t test with Welch’s correction). Quantification of H23 and OVCAR4 colony forming assays treated with 250nM G007-LK or 250nM TDI-012804 normalized to DMSO (n = 3, mean with SD, two-way ANOVA with Tukey correction). (C) Brightfield images of APC-mutant small intestinal organoids treated with DMSO, G007-LK (250nM), or TDI-012804 (250nM) for six days. Scale bar = 800um. (D) Dose response curve of viability of ApcQ1405X small intestinal organoids treated with G007-LK or TDI-012804 for six days measured using AlamarBlue (n = 3, error bars = SEM, see Supplementary Table S5 for 95% CIs). Fluorescence is normalized to DMSO. (E) Organoid viability of ApcQ1405X small intestinal organoids treated with G007-LK (250nM) or TDI-012804 (250nM) for 20 days measured using AlamarBlue (n = 3, mean with SEM, two-way ANOVA with Tukey correction). (F) Dose response curve of viability of ApcQ1405X/KrasG13D/p53Q97X small intestinal organoids treated with G007-LK or TDI-012804 for six days measured using AlamarBlue (n = 3-6, error bars = SEM, see Supplementary Table S5 for 95% CIs). Fluorescence is normalized to DMSO. (G) Dose response curve of viability of ApcQ1405X/KrasG12D/p53Q97X small intestinal organoids treated with G007-LK or TDI-012804 for six days measured using AlamarBlue (n = 3-6, error bars = SEM, see Supplementary Table S5 for 95% CIs). Fluorescence is normalized to DMSO.

We next measured cell proliferation in ApcQ1405X mutant organoids over 6 days of treatment. As expected, G007-LK showed robust suppression of proliferation across all Tnks1 genotypes, as measured by a marked reduction in total organoid area (Fig 5C, fig. S17) and viability/metabolic activity (Alamar blue; Fig. 5D). Likewise, TDI-012804 strongly suppressed Apc^Q1405X^/Tnks1^KO^ organoid proliferation (EC50 59.1nM), while Tnks1 diploid organoids were significantly less sensitive, showing a 19-fold increase in EC50 (1.1uM) (Fig 5D, fig S17). Again, Tnks1^HET^ organoids were sensitive to TNKS2 inhibition, but showed a 3-fold higher EC50 (199.6nM) compared to Tnks1^KO^ organoids (Fig. 5D, fig. S17). To confirm that prolonged treatment with TDI-012804 does not have a cumulative detrimental effect in Tnks1^WT^ cells, we treated organoids for 20 days (refreshing compound every 2 days), measuring metabolic viability and passaging every 4 days. Consistent with the short-term analysis, G007-LK showed robust suppression of organoid growth, while TDI-021804 induced a significant reduction in proliferation in Tnks1^KO^ and Tnks1^HET^ cells, but not Tnks1^WT^ organoids (Fig. 5E). Finally, to determine whether TDI-012804 could suppress proliferation in the context of common CRC oncogenic mutations, we generated Apc^Q1405X^/Kras^G13D^/p53^Q97X^ (AKP-G13D) and Apc^Q1405X^/Kras^G12D^/p53^Q97X^ (AKP-G12D) mutant intestinal organoids with or without Tnks1 loss. Mirroring data in Apc^Q1405X^ mutant cultures, TDI-012804 was selectively toxic to Tnks1^KO^ AKP-G12D and AKP-G13D organoids, compared to Tnks1^WT^ lines (Fig. 5F-G).

## DISCUSSION

WNT hyperactivation is a potent driver in many types of cancer and there is substantial interest in targeting the WNT pathway as a therapeutic strategy. However, multiple studies have identified significant on-target toxicity in the gut and bone from systemic WNT inhibition, highlighting the need for more tumor-selective therapies. Here, we show that loss of TNKS1 expression through cancer-associated chromosome 8p deletions creates a functional dependency on TNKS2 activity, which sensitizes cancer cells to TNKS2-selective inhibition. Further, through rational design, we develop a first-in-class TNKS2-selective small molecule inhibitor and show that this compound can selectively target TNKS1-depleted human cancer cells and murine organoids with APC, KRAS and p53 mutations.

Collateral vulnerability or ‘collateral lethality’ is the emergence of cancer-specific dependency due to loss of large chromosomal regions that delete a redundant ortholog of a key pathway regulator (39, 40). The first example of this concept was described by DePinho and colleagues, who showed that the homozygous loss of the distal region of chromosome 1p (1p36) containing Enolase1 (ENO1), makes glioblastomas selectively dependent on ENO2 (39). Recent development of an ENO2-selective inhibitor supports the therapeutic potential of these strategies for developing cancer-selective treatment approaches (39, 41). Importantly, though the concept of collateral vulnerability was proposed based on homozygous deletion of a redundant ortholog, heterozygous ENO1 cells do show increased sensitivity to ENO2 inhibition (39, 41). Similarly, we show that while TNKS2 blockade is most effective in cells carrying homozygous TNKS1 deletions, heterozygous TNKS1 loss also sensitizes cells to TNKS2-selective inhibition. This is consistent with data from earlier mouse genetic studies showing that while TNKS2 knockout mice are viable and fertile, loss of a single TNKS1 allele in this background is embryonic lethal (31). Together with data described here, these observations suggest that TNKS1 is haploinsufficient for maintaining Tankyrase activity in the context of TNKS2 disruption and expands the potential scope of TNKS2 inhibition to cancers with 8p homozygous and heterozygous loss.

More than 50 Tankyrase inhibitors have been reported in the literature (16-18, 20-23, 32), and in almost all cases, they inhibit both TNKS1 and TNKS2. This is in part due to the high degree of structural similarity in the enzymatic domain of TNKS1 and TNKS2 (Fig. 2A). Through computational modeling, we predicted that the B-loop motif adjacent to the ‘acceptor’ site in TNKS2 is conformationally flexible, assuming structurally ordered ‘open’ and ‘closed’ loop conformations. Specifically, we predicted that presence of a Leucine residue in the TNKS2 B-loop enhances the dynamics of this loop relative to a tyrosine at this position in TNKS1 which possesses stronger intramolecular interactions in the ‘closed’ B-loop conformation. By extending a non-selective inhibitor scaffold through the active site to target the open B-loop, we developed TDI-012804 that selectively inhibits TNKS2. Using cryoEM, we show that binding of TDI-012804 forces movement of this loop to an open conformation. This conformational change is unfavorable in the more rigid TNKS1 B-loop, allowing for selective TNKS2 binding. A similar type of allosteric selectivity has previously been used successfully in the development of mutant KRAS inhibitors, where targeting the S-IIP allosteric pocket in KRAS^G12C^ was able to disrupt functional binding interactions (42). This method of allosteric binding could open many more opportunities to target disease-related proteins that are difficult to target or were previously thought undruggable.

Previous studies have shown that WNT suppression can induce tumor regression, indicating a significant therapeutic potential. However, WNT suppression is limited by dose-limiting toxicities due to the key role of WNT signaling in maintaining healthy tissue (16, 17). While WNT targeted therapies have made little clinical progress, identifying safe and effective strategies to inhibit WNT would have a significant clinical impact. In addition to the direct targeting of WNT-hyperactive cancers, it has been shown that modulating WNT signaling can improve immune-based therapies (43-45). The development of safe WNT therapies would also allow for WNT targeting to be combined with existing treatments, enabling the development of more precise and effective combination therapies. The concept and compounds described here represent a new step towards developing targeted WNT therapies.

## METHODS

### Cell Lines

DLD1 [CCL-221], SW480 [CCL-228], SW1417 [CCL-238], and 293T [CRL-3216] cell lines were purchased from ATCC and cultured in DMEM supplemented with 1% Penicillin-streptomycin and 10% fetal bovine serum. 3T3 cells [CRL-1658] were purchased from ATCC and cultured DMEM supplemented with 1% Penicillin-streptomycin and 10% fetal calf serum. OVCAR-4 cells were purchased from Sigma-Aldrich [SCC258] and cultured in RPMI-1640 supplemented with 1% Penicillin-streptomycin and 10% fetal bovine serum. SW403 cells were purchased from ATCC [CCL-230] and cultured in RPMI-1640 supplemented with 1% Penicillin-streptomycin and 10% fetal bovine serum. PC9 and H23 cells (a gift from Dr. Harold Varmus, Weill Cornell Medicine) were cultured in RPMI-1640 supplemented with 1% Penicillin-streptomycin and 10% fetal bovine serum. HCC1171 cells (a gift from Dr. Gina DeNicola, Moffitt Cancer Center, FL) were cultured in RPMI-1640 supplemented with 1% Penicillin-streptomycin and 10% fetal bovine serum. Sf9 (Spodoptera frugiperda) insect cells were obtained from Expression Systems and cultured in ESF 921 supplemented with penicillin, streptomycin, and Amphotericin B. Cells were tested for mycoplasma using a Mycoplasma PCR Detection Kit (ABM #G238).

### Cloning

The shRNA sequences were cloned into the LT3-GEPIR, SGEP, or SiREP vectors using restriction digest cloning. TNKS2 cDNA was purchased from GenScript (OHu10661) and cloned into the SGEP vector with a 3xFLAG tag (gBlock) using Gibson assembly. L1136Y mutation was introduced to the TNKS2 cDNA using overlap PCR, and hygromycin resistance was introduced using restriction digest cloning. TOPFlash sequences were cloned into the SGEN vector using Gibson assembly. AXIN1-GFP reporter sequence was cloned into the SGEN vector, and the Kozak sequence was replaced using Gibson assembly. The SAM-PARP coding region of TNKS2 (amino acids 850 -1166) was PCR amplified and cloned into a pFastbac expression vector featuring a N-terminal twin-strep-MBP tag utilizing Gibson assembly. Site directed mutagenesis to generate the L1136Y mutant was performed through Gibson assembly. sgRNAs were cloned into the LRT2B vector or SFFV. NLS-mScarlet vector. shRNA sequences, sgRNA sequences, and cloning primers are listed in Supplementary Table 4.

### Compound Synthesis

Compounds were synthesized by WuXi AppTec. Chemical composition and purity was validated through 1H-NMR, 13C-NMR, LCMS, and HRMS (See Supplementary Methods).

### Lentiviral transduction

HEK-293T cells were seeded to 90% confluency one day prior to transfection. The cells were co-transfected with 5ug of plasmid DNA, 2.5ug PAX2 vector, and 1.25ug VSVg vector. shRNA plasmids were additionally transfected with 1.25ug pcSUPER-shPasha plasmid. Vectors were combined in 150ul DMEM with 30ul PEI and incubated at RT for 10 minutes. Transfection mix was incubated with the 293Ts overnight. Cells were then transferred to collection media, and lentiviral supernatants were collected at 24 and 48h following the change to collection media. Remaining HEK-293T cells were removed through centrifugation. Target cells were seeded to 30-50% confluency. After 24h, the target cells were incubated overnight with diluted lentiviral supernatant mixed with 8ug/ mL polybrene. After 24h, the viral media was replaced with culture media. Antibiotic selection was initiated 48h following transduction.

### Generation of TNKS1 KO cell lines

DLD1 cells were transduced with Cas9-p2a-puro and selected with 2ug/mL puromycin. Cas9-expressing cells were then transduced with an sgRNA sequence targeting TNKS1 (LRT2B or NLS-mScarlet) (sgRNA sequences listed in Supplementary Table 4) and selected with 5ug/mL blasticidin or 400ug/mL neomycin. Transduced cells were plated as single cells by limiting dilution to identify clonal lines.

### Intestinal organoid isolation and culture

Mouse small intestinal organoids were isolated as described previously (46). Organoids were plated in basal medium (Advanced DMEM/F12 containing penicillin/streptomycin, glutamine, and HEPES) containing EGF [40 ng/ml], LDN [50nM LDN], RSPO1 [5% conditioned media], and Fungin [10ug/ml]. Organoids were maintained by passaging every 2-3 days. To passage the organoids, the Matrigel containing organoids was collected in PBS. The organoids were mechanically separated using a P1000 pipette and washed with additional PBS. The organoids were spun at 1200 rpm for 4 minutes. The PBS was aspirated, and the organoid pellet was resuspended in Matrigel. After Matrigel polymerization (5 mins at RT, 5 mins at 37C), organoids were cultured in ENR media (basal medium containing EGF [40 ng/ml], LDN [50nM], RSPO1 [5% conditioned media]).

### Crypt survival

Mice were treated with doxycycline for 2 or 3 weeks. Small intestinal crypts were isolated as described above and plated in multiple wells of a 48-well plate (40ul Matrigel / well). Organoids were plated in ENR media (described above) containing 0.5ug/mL doxycycline when indicated. ENR + dox media was replaced on day 2. Viable organoids were counted 12 hours and 4 days after plating using a brightfield microscope.

### Organoid transfection

Prior to transfection, organoids were cultured in media containing CHIR99021 (5 uM) and Y-27632 (10 uM) for two days. Organoids were dissociated to single cells by incubating in 200uL TrypLE at 37C for 5 minutes, centrifuged, and resuspended in 600uL culture media. 2ug of DNA was mixed with 4uL Lipofectamine 2000 in 200uL Opti-MEM, and the transfection mix was combined with the organoid suspension in a 24-well plate. The plate was then centrifuged at 600g for 60 minutes at 32C. Following centrifugation, the plate was incubated at 37C for an additional 4 hours. The organoids were pelleted and replated in Matrigel in media containing CHIR99021 (5 uM) and Y-27632 (10 uM) for 2 days to facilitate organoid recovery. To select for APC mutations, RSPO1 and LDN were removed from the media 2 days after transfection. To select for p53 mutations, the organoids were treated with Nutlin (10uM).

### Organoid proliferation and viability

Organoids were plated at low density in 40ul of Matrigel in 24-well plates and cultured in EN media (Basal media + EGF + LDN, described above). Organoids were grown continually for 6 days in media containing DMSO or inhibitor compounds, with compounds replenished by changing media every two days. Organoid area was quantified using the Sartorius Incucyte S3. After 6 days of compound treatment, alamarBlue Cell Viability Reagent (ThermoFisher #DAL1100) was added to each well at a 1:10 dilution and incubated at 37C for 4 hours. 100ul of media was transferred to a 96-well plate and fluorescence intensity was quantified using a BMG Labtech Fluostar Omega microplate reader. For serial passaging experiments, organoids were plated at high density in 120ul of Matrigel each in 2 wells per condition of a 12-well plate in EN media containing DMSO or inhibitor compound. Media was changed every 2 days. Every 4 days, 100ul of alamarBlue was added to one well per condition and quantified as above. The second well was passaged 1:2 into the same condition in a new 12-well plate.

### Western blot

700,000 cells were seeded in each well of a 6-well plate and incubated for 24 hours for the cells to adhere. Compounds were then added to the cells and incubated for 24 hours. To isolate protein, cells were washed in PBS and lysed in 200ul RIPA buffer on ice for 20 minutes. Lysates were centrifuged at 15,000 rpm for 10 minutes and the protein supernatant was stored at -80C. 20ug of protein was run per sample for western blot analysis. Antibodies used were anti-AXIN1 [Cell Signaling Technology, #2087; 1:1000], anti-TNKS1/2 [Santa Cruz Biotechnology, #sc-365897; 1:1000], anti-GFP [Cell Signaling Technology, #2956; 1:1000], and anti-beta Actin-HRP [Abcam, #ab49900; 1:10,000]. When blotting for TNKS1/2, cells were incubated in 1uM XAV939 for 24h prior to cell lysis unless otherwise indicated. Protein abundance was quantified using Image Lab software.

### Biochemical PARylation assays

TNKS1 and TNKS2 enzymatic activity was measured using the BPS Bioscience TNKS1 (#80573) and TNKS2 (#80572) Histone Ribosylation assay kit according to the manufacturer’s instructions. Chemiluminescence was measured using a BMG Labtech Fluostar Omega microplate reader.

### AXIN1-GFP flow cytometry

GFP reporter cell lines were seeded 50,000 cells per well in 24-well plates. The cells were incubated for 24 hours to allow the cells to adhere prior to starting compound treatments. After adhering, the media was replaced with culture media containing inhibitor compounds. The compounds were diluted in DMSO at 1000x final concentration, then diluted 1:1000 in cell culture media. The cells were treated for 24 hours. Following this incubation, the cells were trypsinized in 100ul at 37C for 5 minutes, then resuspended in 300ul culture media containing DAPI. 300ul of the resuspension was transferred to a round-bottom 96-well plate and fluorescence was measured on a ThermoFisher Attune NxT flow cytometer. Results were analyzed using Flowjo software.

### TOPflash assay

TOPflash expressing cell lines were seeded 15,000 cells per well in clear-bottom black 96-well plates. The plates were incubated for 24 hours to allow the cells to adhere. Following 24 hours, the compounds were serially diluted in cell culture media using a multichannel pipette and replaced the media in the cell culture plates. Following 48 hours of compound treatment, firefly and renilla luciferase were analyzed using the BPS Bioscience Dual Luciferase (Firefly-Renilla) Assay System (#60683-2). The assay was performed according to the manufacturer’s instructions, and luminescence was recorded using a BMG Labtech Fluostar Omega microplate reader.

### Colony Forming Assays

Cells were seeded at 1000 cells per well into 12-well plates and incubated for 24h for cells to adhere. Compounds were added to the media and were replenished every two days. Once colonies formed (about 16 days after compounds were added), the media was removed, and the cells were fixed in 4% paraformaldehyde for one hour at RT on an orbital shaker. The PFA was removed, and the fixed cells were stained in a 10% Giemsa solution in PBS overnight on an orbital shaker. After staining, the Giemsa solution was removed, the plates were washed in water, and left to dry overnight. Once dry, the plates were imaged using an Epson Perfection V550 Photo scanner. To quantify cell growth, 1mL of 10% acetic acid was added to each well. The plate was shaken for 20 minutes, and 100ul of the acetic acid solution was transferred to a 96-well plate. Absorbance was quantified using a BMG Labtech Fluostar Omega microplate reader.

### RNA isolation and RNA-seq

One six-well of organoids was plated per condition into EN media (basal medium containing EGF [40 ng/ml] and LDN [50nM LDN]) containing DMSO, 250nM G007-LK, or 500nM TDI-012804. Organoids were cultured for 3 days, with media replenished on the second day. The organoids were collected in 1mL of TRIzol (Thermo Fisher Scientific, #15596018) and RNA was extracted following the manufacturer’s protocol. DNA was removed from the isolated RNA by treating with DNase1 for 15 minutes followed by column purification with the Qiagen RNeasy kit (Qiagen #74106). RNA sequencing was performed by the Genomics Core Laboratory at Weill Cornell Medicine: the RNA quality was confirmed using a 2100 Bioanalyzer (Agilent technologies), the RNA library was prepared using TruSeq Stranded mRNA Sample Library Preparation Kit (Illumina), and RNA-seq was performed on an Illumina NovaSeq 6000 with paired-end 2×100 cycles.

### RNAseq analysis

Transcript abundance was estimated using Kallisto (47), aligned to the GRCm38 mouse reference genome. Transcript per million (TPM) data was reported for each gene after mapping gene symbols to ensemble IDs using R packages, “tximport”, tximportData”, “ensembldb”, and “EnsDb. Mmusculus.v79”. Differential gene expression was estimated using DESeq2 (48). For data visualization and gene ranking, log fold changes were adjusted using lfcShrink in DESeq2, to minimize the effect size of poorly expressed genes. GSEA analysis (v3.0) was performed on pre-ranked gene sets from differential expression between control and treated groups. We used R (v3.6.1) and R Studio (v1.2.1335) to create all visualizations, perform hierarchical clustering and principal component analysis. Visualizations of RNAseq data were produced using Enhanced Volcano (volcano plots), pheatmap (heatmaps), and ggplot2 (dotplots).

### Lgr5 ISH

Freshly cut 5-micron paraffin sections were stained using RNAscope 2.5 LS Red kit (ACD, cat#322150) and Bond Polymer Refine Red Detection kit (Leica, cat#DS9390) on Leica Bond RX instrument following routine manufacturer protocol ACD 2.5 Red. RNAscope 2.5 LS probes for Ms-LGR5 (ACD, cat# 312178). DapB-negative control (ACD, cat#312038) were used with hybridization at 42C for 2 hours. The sections were pre-treated with Leica Bond ER2 Buffer for 20 min at 95C and Protease III (ACD, cat#322102) for 20 min at 40C. After staining the sections were counterstained with Hematoxylin and 10ug/ml DAPI for 10 min and mounted with Mowiol mounting media.

### Sf9 insect cell protein expression

WT TNKS, TNKS2 SAM-PARP and associated mutants were expressed in Sf9 insect cells. Viral bacmids were generated using Tn7 transposition in chemically competent DH10Bac E. coli cells (Thermo Scientific #10361012). Bacmids were purified using a Bacmid DNA miniprep kit (Zymo Research #D4049) and transfected into Sf9 cells using Cellfectin II (Thermo Scientific #10362100) to generate recombinant baculoviruses. Proteins were expressed in 800 ml of Sf9 cells that were infected with the P2 amplified baculovirus at a cell density of 3×106 cells per ml. Infected Sf9 cells were incubated at 28ºC with mild shaking (130 rpm) until viability dropped to approximately 80% (typically 72 hours post infection). Cell pellets were collected via centrifugation at 500 g for 20 minutes and resuspended in lysis buffer (500 mM NaCl, 20 mM HEPES pH 7.5, 2 mM MgCl2, 5% v/v glycerol and 2 mM 2-mercaptoethanol which was supplemented with cOmplete EDTA free protease inhibitor cocktail (Roche #11697498001) and DNase I (Worthington # LS006342).

### Protein purification

Cells, resuspended in lysis buffer, were lysed by sonication and centrifuged at 48,000 g to remove insoluble cell debris. Lysates were filtered with a 5-uM filter and loaded onto a 5-ml StrepTrap XT affinity column (Cytiva) using an AKTA PURE. Following loading of the lysate, the column was washed with at least 5 column volumes of wash buffer (lysis buffer without additional supplementation). Proteins were eluted by applying 5 column volumes of elution buffer consisting of the wash buffer supplemented with 50 mM Biotin (IBA #2-1016-002). The eluted fractions, containing tankyrase, were dialyzed overnight against 500 mM NaCl, 20 mM HEPES pH 7.5, 2 mM MgCl2, 5% v/v glycerol, 0.05 % tween-20 and 2 mM 2-mercaptoethanol without the addition of preScission protease. The dialyzed samples were concentrated, supplemented to 20% v/v glycerol, flash-frozen in liquid nitrogen and stored at -80ºC.

#### Cryo-EM grid preparation and data collection

3.5 uL of purified protein, at approximately 20 μM, in a buffer containing 150 mM NaCl, 20 mM HEPES pH 7.5, 5 % v/v glycerol, 0.05 % tween-20 and 2 mM 2-mercaptoethanol were applied onto Quantifoil Au R 1.2/1.3 400 mesh graphene oxide coated grids for 45 seconds in a humidity-controlled vitrobbot Mark IV plunge freezer (Thermo Scientific).

Humidity was set to 95 % and temperature was set to 10ºC. After 45 seconds, 4 μL of a 20 mM HEPES pH 7.5 solution was applied to the grid and removed two times to lower the salt concentration, prior to blotting and vitrification. For all three datasets, grids were imaged using a Titan KRIOS microscope (Thermo Scientific) operated at 300 keV and recorded with a Gatan K3 camera. Micrograph movies were recorded in super-resolution mode (nominal pixel size 0.826 Å), using a total dose of ∼55 electron per Å2, under an applied defocus ranging from -0.8 to -3.0 uM.

### Cryo-EM image processing

For the three datasets, all processing was conducted using cryosparc (v4.4) (49). Movies were motion-corrected and binned 2x prior to patch CTF refinement. On-the-fly image processing utilized the blob picker particle picking algorithm alongside iterative 2D-classification to identify good segments of the TNKS filaments. Subsequent particle picking was performed using the filament tracer job using the previously selected particles’ class averages as templates. The initial volume was reconstructed using cryosparc’s ab-initio reconstruction job in which the output was utilized to estimate the helical symmetry parameters using the symmetry search function. Helical reconstructions were conducted using the helical refinement job in which the helical parameters converged to a helical twist of -52.4º and rise of 13.6 Å. The consensus class was further optimized through iterative global and local CTF refinements as well as through reference-based motion correction as implemented in cryosparc v4.4. Prior to the final refinement, the consensus class was symmetry expanded based on the helical and point group D1 symmetry and the exposure optics groups were expanded according to the beam image shift grouping. Subsequently, per-particle CTF refinement was ran prior to local refinement in C1 to yield the final consensus reconstruction. Local refinement jobs utilizing masks around the central helical unit as well as the most visually well-resolved point group D1 symmetry unit were ran to generate the maps used for model building and refinement. The respective maps’ local resolution was calculated using Bsoft and subsequently applied to generate a locally filtered map volume for experimental interpretation and model building. For all reconstructions, the global resolution denoted in supplementary pipelines was estimated based on the gold-standard Fourier shell correlation (FSC) 0.143 criterion between two independently refined half maps. Data collection parameters and model statistics are shown in Supplementary Table 2. The processing steps for the three datasets are described in Supplementary figures 10-13.

### Model building and refinement

The starting model for the TNKS2 SAM-PARP protomer was generated by manually fitting PDB 8ALY into the central helical unit local refinement using ChimeraX (v 1.6) and removing all chains other than chain D, corresponding to one of the more well-resolved regions of the map. This fitted protomer was re-built in Coot and Isolde followed by real-space refined in Phenix in an iterative fashion. During this process, the ligand if present, was fitted into the map and refined using a restraints file generated by Phenix eLBow. The final optimized protomer model was symmetry expanded in ChimeraX using either D1 symmetry alone or D1 and helical symmetry to generate models that fitted the local refinement maps and subsequently real-space refined. The respective final models were validated using MolProbity, as implemented in Phenix, in which the output validation statistics are shown in Supplementary Table 2.

### Computational model

All calculations were performed using the Schrodinger Suite (version 2019-4 to 2023-3). Free energy calculations were performed using the Schrodinger FEP+ method (Ref: Journal of the American Chemical Society (2015), 137 (7), 2695-2703CODEN: JACSAT; ISSN:0002-7863.) with TNKS2 structure (PDB: 3MHK). The calculations were run for 10ns or until they reached convergence. The ensemble docking model (Glide SP docking) was set up with 18 grids prepared from TNKS2 crystal structures with complete B-loop (3MHK, 4HYF,4PML, 4PNL,4PNM, 4PNN, 4PNQ, 4PNS, 4PNT, 4TJU, 4TJW, 4TJY, 4TK0, 4TK5, 4TKF, 4TKG, 4TKI), together with 2 grids that were generated from the FEP snapshot where the B-loop opened during the simulation. All proteins were prepared using the Protein Preparation Wizard with the default setting. Ligands were prepared using LigPrep.

### Animal Studies

All mouse treatments and experiments were approved by the Institutional Animal Care and Use Committee (IACUC) at Weill Cornell Medicine under protocol 2014-0038. Tnks1 KO mice were produced by the Mouse Genetics Core Facility at the Memorial Sloan Kettering Cancer Center through zygote electroporation of RNPs in C57Bl/6 mice. Mice were genotyped for Col1a1, R26, TGM-shRen.713, TGM-shTnks2.3004, TGM-shTnks1/2.13-30, TGM-shTnks1/2.33-13, Tnks, Kras, and Apc^Q1405X^ (Genotyping primers listed in Supplementary Table 4). For crypt survival experiments, mice were administered doxycycline via food pellets (200mg/kg) for 2 or 3 weeks at 6-9 weeks of age. For Lgr5 ISH, mice were administered doxycycline via food pellets (200mg/kg) for 3 weeks at 6-9 weeks of age.

## Acknowledgements

We thank members of the Dow lab for advice and comments on preparation of the manuscript. The authors gratefully acknowledge the support to the project generously provided by the Sanders Tri-Institutional Therapeutics Discovery Institute (TDI), a 501(c)(3) organization. TDI receives financial support from Takeda Pharmaceutical Company, TDI’s parent institutes (Memorial Sloan Kettering Cancer Center, The Rockefeller University and Weill Cornell Medicine) and from a generous contribution from Mr. Lewis Sanders and other philanthropic sources. We thank Jason de la Cruz at the Memorial Sloan Kettering Cancer Center (MSKCC) Richard Rifkind Center for cryo-EM assistance with data collection and the MSKCC High-Performance Computing (HPC) group for assistance with cryo-EM data processing. The content is solely the responsibility of the authors and does not necessarily represent the official views of the NIH.

## Authors contributions

JZ designed and performed experiments, analyzed data, and wrote the paper. BM, EFG, SS, RL, DJH, TK, JG, NL, MF, EMS, EG, SG, AK, BL, HA, and YF performed experiments and/or analyzed data. TT, RL, and RKH analyzed data and supervised experimental work. LED designed and supervised experiments, analyzed data and wrote the paper.

## Data and materials availability

The cryo-EM density maps and atomic coordinates have been deposited in the EMDataBank and PDB as follows: map1 (EMD-43738, PDB 8W23), map2 (EMD-43739, PDB 8W25), map3 (EMD-43740, PDB 8W27), map4 (EMD-43471, PDB 8W28), map5 (EMD-43759, PDB 8W2U) and map6 (EMD-43758, PDB 8W2T). The atomic model used for initial model building and analysis is available from the Protein Data Bank under the accession code 8ALY. Raw RNA sequencing data is deposited at the sequence read archive (SRA) under accession PRJNA1079707.

## REFERENCES

1. L. E. Dow et al., Apc Restoration Promotes Cellular Differentiation and Reestablishes Crypt Homeostasis in Colorectal Cancer. Cell 161, 1539–1552 (2015).

2. K. P. O’Rourke et al., Transplantation of engineered organoids enables rapid generation of metastatic mouse models of colorectal cancer. Nat Biotechnol 35, 577–582 (2017).

3. E. M. Schatoff et al., Distinct Colorectal Cancer-Associated APC Mutations Dictate Response to Tankyrase Inhibition. Cancer Discov 9, 1358–1371 (2019).

4. T. Han et al., Lineage Reversion Drives WNT Independence in Intestinal Cancer. Cancer Discov 10, 1590–1609 (2020).

5. E. E. Storm et al., Targeting PTPRK-RSPO3 colon tumours promotes differentiation and loss of stem-cell function. Nature 529, 97–100 (2016).

6. B. Madan et al., Wnt addiction of genetically defined cancers reversed by PORCN inhibition. Oncogene 35, 2197–2207 (2016).

7. A. Scholer-Dahirel et al., Maintenance of adenomatous polyposis coli (APC)-mutant colorectal cancer is dependent on Wnt/beta-catenin signaling. Proc Natl Acad Sci U S A 108, 17135–17140 (2011).

8. M. C. Faux et al., Restoration of full-length adenomatous polyposis coli (APC) protein in a colon cancer cell line enhances cell adhesion. J Cell Sci 117, 427–439 (2004).

9. P. Polakis, The many ways of Wnt in cancer. Curr Opin Genet Dev 17, 45–51 (2007).

10. O. Arqués et al., Tankyrase Inhibition Blocks Wnt/β-Catenin Pathway and Reverts Resistance to PI3K and AKT Inhibitors in the Treatment of Colorectal Cancer. Clin Cancer Res 22, 644–656 (2016).

11. C. Y. Fong et al., BET inhibitor resistance emerges from leukaemia stem cells. Nature 525, 538–542 (2015).

12. P. Rathert et al., Transcriptional plasticity promotes primary and acquired resistance to BET inhibition. Nature 525, 543–547 (2015).

13. M. Schoumacher et al., Inhibiting Tankyrases sensitizes KRAS-mutant cancer cells to MEK inhibitors via FGFR2 feedback signaling. Cancer Res 74, 3294–3305 (2014).

14. S. Spranger, R. Bao, T. F. Gajewski, Melanoma-intrinsic β-catenin signalling prevents anti-tumour immunity. Nature 523, 231–235 (2015).

15. M. Ruiz de Galarreta et al., β-Catenin Activation Promotes Immune Escape and Resistance to Anti-PD-1 Therapy in Hepatocellular Carcinoma. Cancer Discov 9, 1124–1141 (2019).

16. T. Lau et al., A novel tankyrase small-molecule inhibitor suppresses APC mutation-driven colorectal tumor growth. Cancer Res 73, 3132–3144 (2013).

17. Y. Zhong et al., Tankyrase Inhibition Causes Reversible Intestinal Toxicity in Mice with a Therapeutic Index < 1. Toxicol Pathol 44, 267–278 (2016).

18. S. M. Huang et al., Tankyrase inhibition stabilizes axin and antagonizes Wnt signalling. Nature 461, 614–620 (2009).

19. W. Wang et al., Tankyrase Inhibitors Target YAP by Stabilizing Angiomotin Family Proteins. Cell Rep 13, 524–532 (2015).

20. B. Chen et al., Small molecule-mediated disruption of Wnt-dependent signaling in tissue regeneration and cancer. Nat Chem Biol 5, 100–107 (2009).

21. J. Waaler et al., A novel tankyrase inhibitor decreases canonical Wnt signaling in colon carcinoma cells and reduces tumor growth in conditional APC mutant mice. Cancer Res 72, 2822–2832 (2012).

22. M. D. Shultz et al., Identification of NVP-TNKS656: the use of structure-efficiency relationships to generate a highly potent, selective, and orally active tankyrase inhibitor. J Med Chem 56, 6495–6511 (2013).

23. C. C. Mehta, H. G. Bhatt, Tankyrase inhibitors as antitumor agents: a patent update (2013 - 2020). Expert Opin Ther Pat 31, 645–661 (2021).

24. Y. Cai et al., Loss of Chromosome 8p Governs Tumor Progression and Drug Response by Altering Lipid Metabolism. Cancer Cell 29, 751–766 (2016).

25. C. E. Gustafson et al., Functional evidence for a colorectal cancer tumor suppressor gene at chromosome 8p22-23 by monochromosome transfer. Cancer Res 56, 5238–5245 (1996).

26. M. L. Yaremko, W. M. Recant, C. A. Westbrook, Loss of heterozygosity from the short arm of chromosome 8 is an early event in breast cancers. Genes Chromosomes Cancer 13, 186–191 (1995).

27. I. I. Wistuba et al., Allelic losses at chromosome 8p21-23 are early and frequent events in the pathogenesis of lung cancer. Cancer Res 59, 1973–1979 (1999).

28. W. Xue et al., A cluster of cooperating tumor-suppressor gene candidates in chromosomal deletions. Proc Natl Acad Sci U S A 109, 8212–8217 (2012).

29. I. T. P.-C. A. o. W. G. Consortium, Pan-cancer analysis of whole genomes. Nature 578, 82–93 (2020).

30. M. Foronda et al., Tankyrase inhibition sensitizes cells to CDK4 blockade. PLoS One 14, e0226645 (2019).

31. Y. J. Chiang et al., Tankyrase 1 and tankyrase 2 are essential but redundant for mouse embryonic development. PLoS One 3, e2639 (2008).

32. M. Yu, Y. Yang, M. Sykes, S. Wang, Small-Molecule Inhibitors of Zimmerman et al, bioRxiv preprint Tankyrases as Prospective Therapeutics for Cancer. J Med Chem 65, 5244–5273 (2022).

33. A. Nathubhai et al., Highly Potent and Isoform Selective Dual Site Binding Tankyrase/Wnt Signaling Inhibitors That Increase Cellular Glucose Uptake and Have Antiproliferative Activity. J Med Chem 60, 814–820 (2017).

34. S. Tomassi et al., From PARP1 to TNKS2 Inhibition: A Structure-Based Approach. ACS Med Chem Lett 11, 862–868 (2020).

35. A. Nathubhai et al., Structure-activity relationships of 2-arylquinazolin-4-ones as highly selective and potent inhibitors of the tankyrases. Eur J Med Chem 118, 316–327 (2016).

36. N. Pillay et al., Structural basis of tankyrase activation by polymerization. Nature 612, 162–169 (2022).

37. L. Mariotti et al., Tankyrase Requires SAM Domain-Dependent Polymerization to Support Wnt-β-Catenin Signaling. Mol Cell 63, 498–513 (2016).

38. L. Mygland et al., Identification of response signatures for tankyrase inhibitor treatment in tumor cell lines. iScience 24, 102807 (2021).

39. F. L. Muller et al., Passenger deletions generate therapeutic vulnerabilities in cancer. Nature 488, 337–342 (2012).

40. F. L. Muller, E. A. Aquilanti, R. A. DePinho, Collateral Lethality: A new therapeutic strategy in oncology. Trends Cancer 1, 161–173 (2015).

41. Y. H. Lin et al., An enolase inhibitor for the targeted treatment of ENO1-deleted cancers. Nat Metab 2, 1413–1426 (2020).

42. J. M. Ostrem, U. Peters, M. L. Sos, J. A. Wells, K. M. Shokat, K-Ras(G12C) inhibitors allosterically control GTP affinity and effector interactions. Nature 503, 548–551 (2013).

43. J. Waaler et al., Tankyrase inhibition sensitizes melanoma to PD-1 immune checkpoint blockade in syngeneic mouse models. Commun Biol 3, 196 (2020).

44. N. C. DeVito et al., Pharmacological Wnt ligand inhibition overcomes key tumor-mediated resistance pathways to anti-PD-1 immunotherapy. Cell Rep 35, 109071 (2021).

45. T. Huang et al., Wnt Inhibition Sensitizes PD-L1 Blockade Therapy by Overcoming Bone Marrow-Derived Myofibroblasts-Mediated Immune Resistance in Tumors. Front Immunol 12, 619209 (2021).

46. K. P. O’Rourke, S. Ackerman, L. E. Dow, S. W. Lowe, Isolation, Culture, and Maintenance of Mouse Intestinal Stem Cells. Bio Protoc 6, (2016).

47. N. L. Bray, H. Pimentel, P. Melsted, L. Pachter, Near-optimal probabilistic RNA-seq quantification. Nat Biotechnol 34, 525–527 (2016).

48. M. I. Love, W. Huber, S. Anders, Moderated estimation of fold change and dispersion for RNA-seq data with DESeq2. Genome Biol 15, 550 (2014).

49. A. Punjani, J. L. Rubinstein, D. J. Fleet, M. A. Brubaker, cryoSPARC: algorithms for rapid unsupervised cryo-EM structure determination. Nat Methods 14, 290–296 (2017).

