## Supplementary_Figures for "A potent and selective TNKS2 inhibitor for tumor-selective WNT suppression"

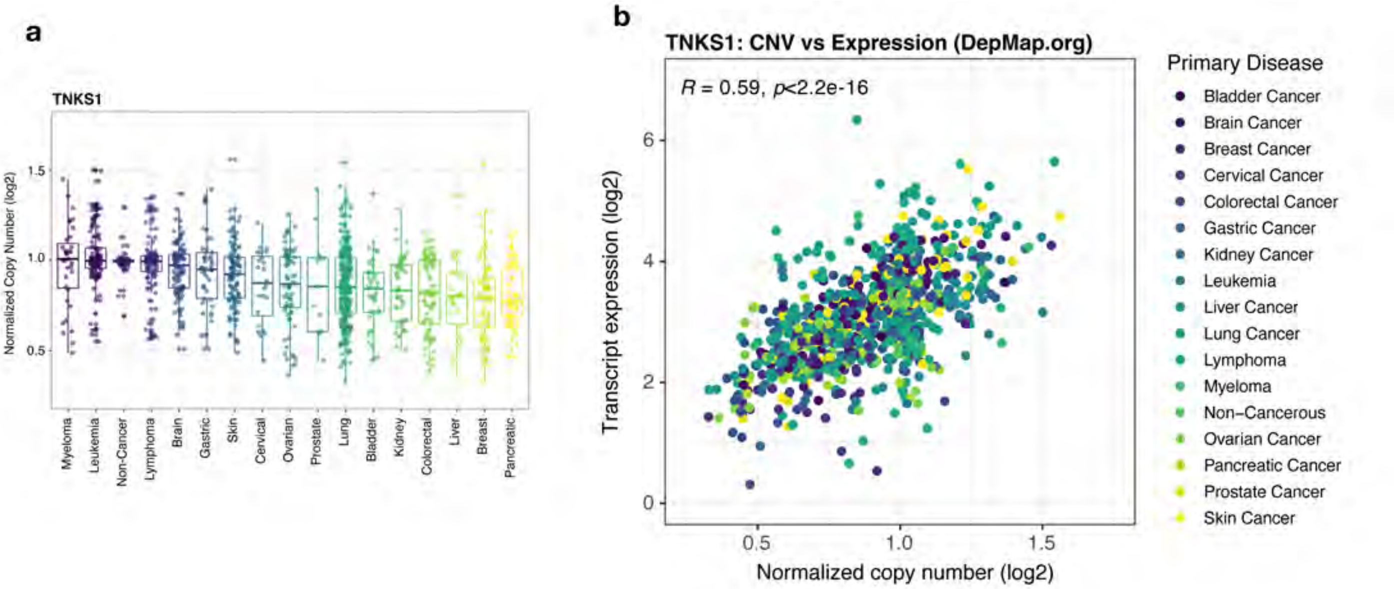


**Figure S1: TNKS1 expression in 8p-deleted cancers**

**a.** Normalized TNKS1 copy number across cancer cell lines. **b.** Scatter plot showing TNKS1 copy number (log2) vs transcript expression across cancer cell lines. *Primary data from DepMap.or*g

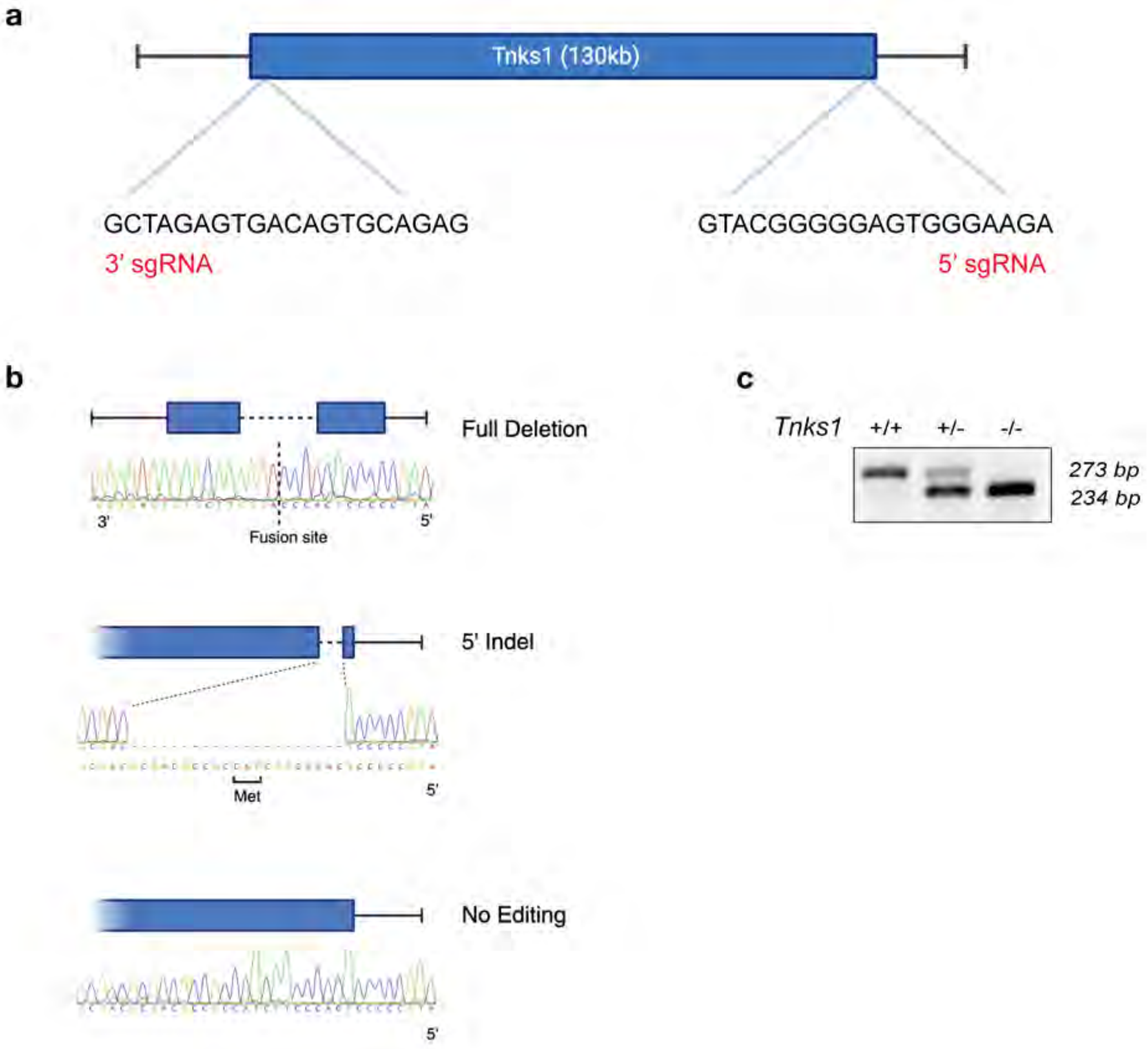


**Figure S2: Generation and validation of a Tnks1 KO mouse**

**a**. Schematic showing dual sgRNAs targeting 5' and 3' ends of *Tnks1* gene. Created with Biorender.com **b**. Examples of modified alleles in founder animals, showing full gene deletion, isolated indels, or no editing. Most full deletion animals showed the same fusion sequence due to microhomology within guide sequences. Created with Biorender.com **c**. PCR genotyping of Tnks1 KO mice. Upper band is amplification of 5' terminal. Lower band is amplification of genomic fusion product following KO.

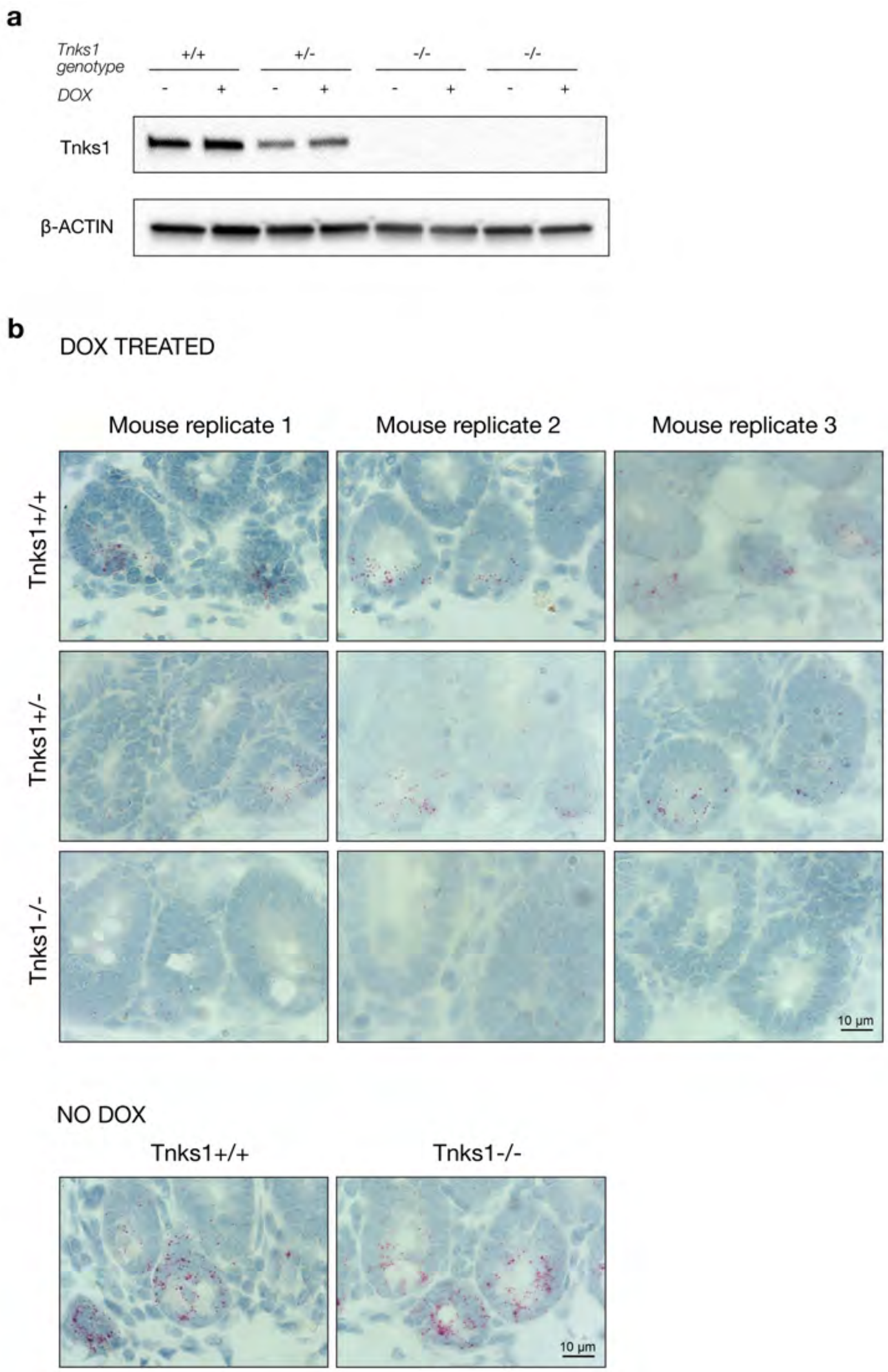


**Figure S3: Selective loss of Lgr5 expression in small intestinal crypts**

**a.** Western blot showing loss of TNKS1 protein in small intestinal organoids derived from *Tnks1*$^{WT}$, *Tnks1*$^{Het}$, or *Tnks1*$^{KO}$ mice. **b.** In situ hybridization of *Lgr5* mRNA expression in the small intestine of *shTnks2/Tnks1*$^{WT}$, *shTnks2/Tnks1*$^{Het}$, and *shTnks2/Tnks1*$^{KO}$ mice fed doxycycline chow (200mg/kg) for 3 weeks or maintained on regular chow (no dox; no shRNA induction). Each image shows a representative section from an independent mouse.

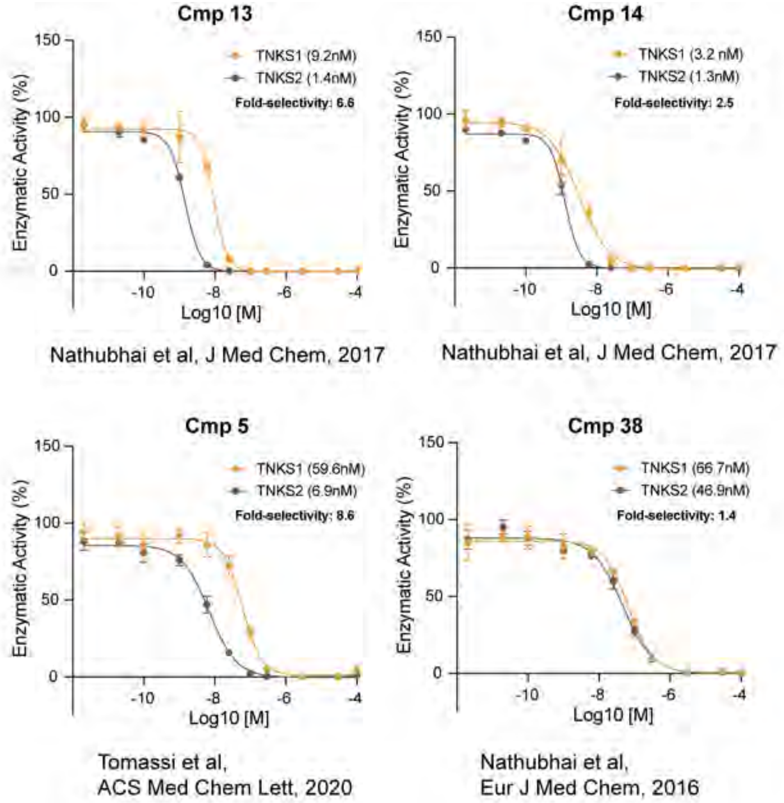


**Figure S4. TNKS1/2 enzymatic inhibition with published TNKS2-selective inhibitors**
Dose response curves from in vitro PARylation assays measuring the enzymatic activity of TNKS and TNKS2 incubated with published TNKS2 inhibitors (n = 2, error bars = SEM). Enzymatic activity is normalized to DMSO.

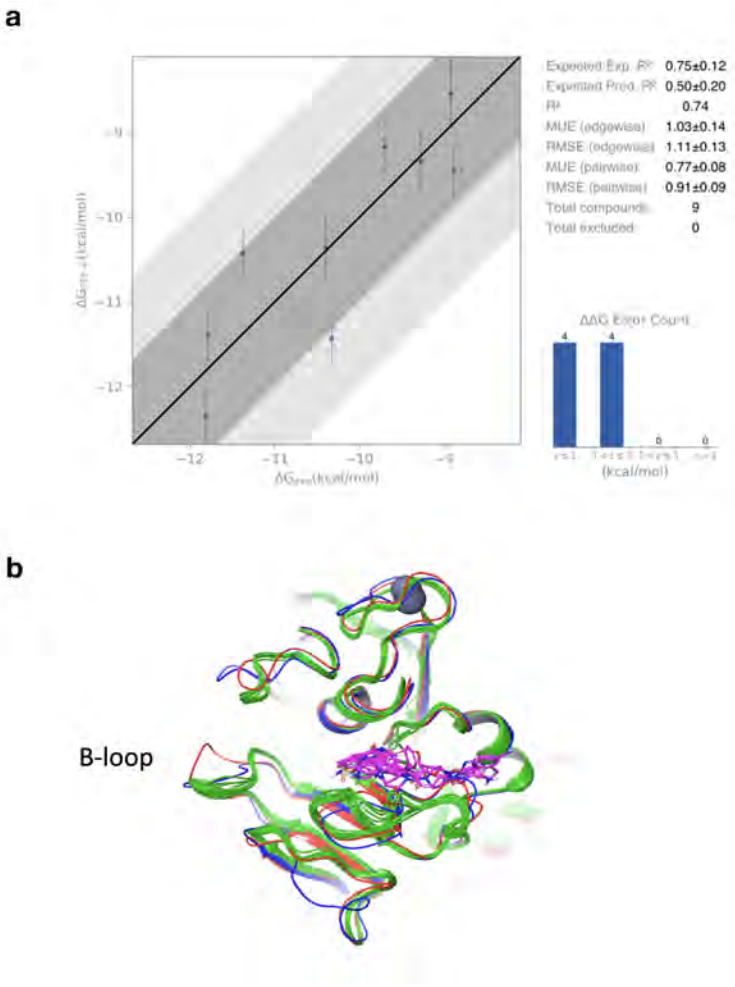


**Figure S5: Docking model of TNKS2 compounds**

**a.** FEP validation plot **b.** Aligned grids used for TNKS2 ensemble docking. Grids in green ribbons are TNKS2 crystal structures. Grids shown in red and blue ribbons are modeled structures from FEP calculation.

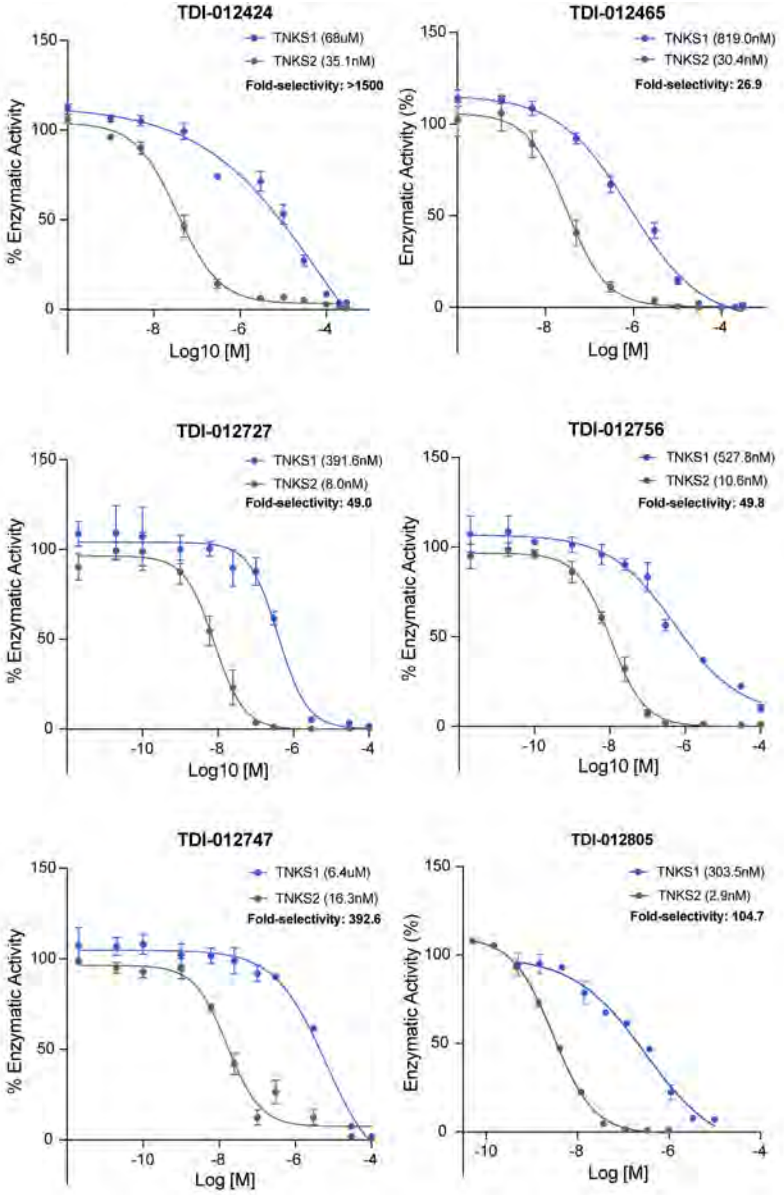


**Figure S6. TNKS1/2 enzymatic inhibition with potent and selective hit compounds**

Dose response curves from in vitro PARylation assays measuring the enzymatic activity of TNKS1 and TNKS2 incubated with indicated compounds (n = 2, error bars = SEM). Enzymatic activity is normalized to DMSO.

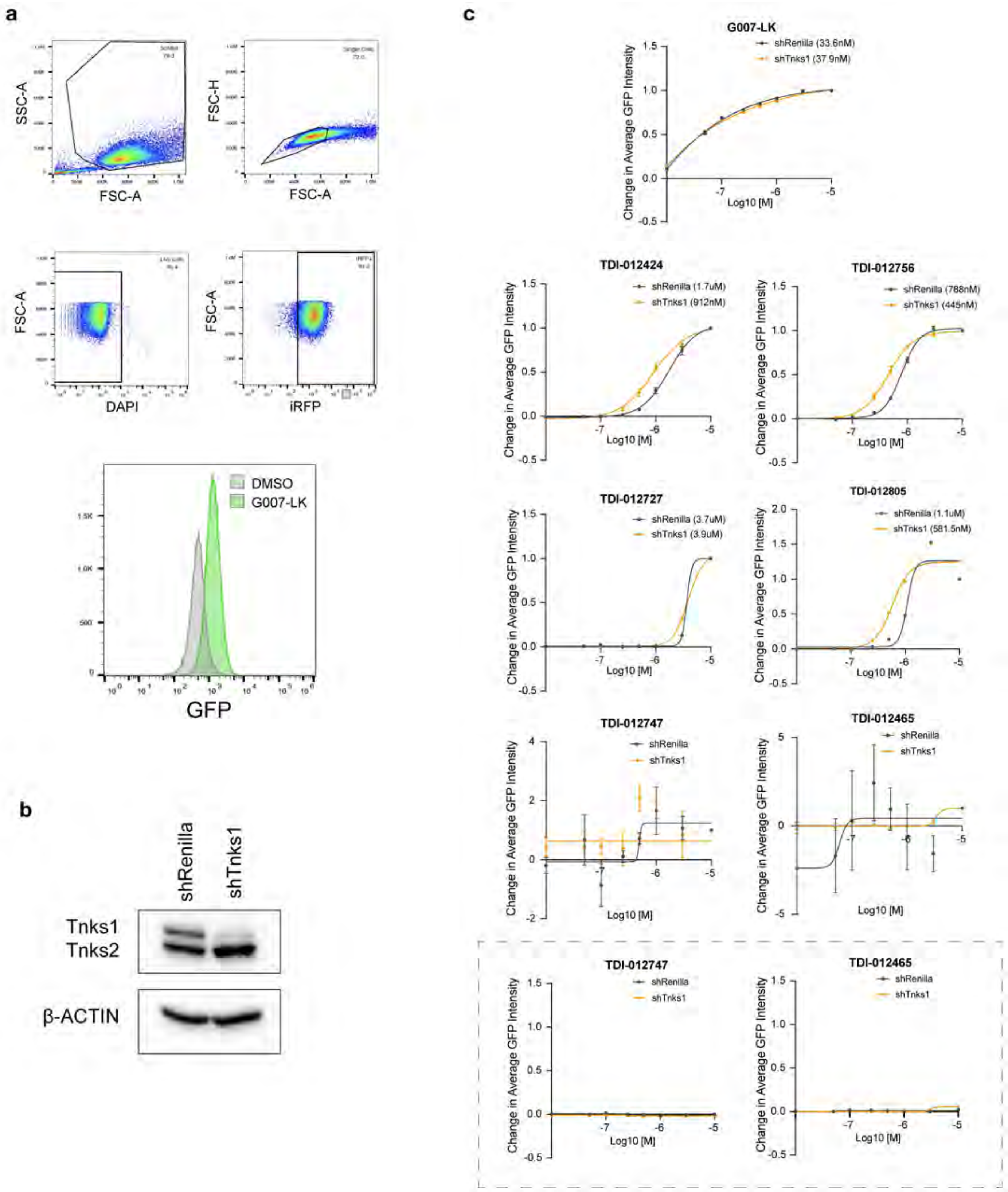


**Figure S7. AXIN1-GFP reporter cell line to quantify AXIN1 stabilization**

**a.** Flow cytometry gating strategy to determine average GFP intensity in AXIN1-GFP reporter cells. **b.** Western blot showing a decrease in Tnks protein levels in AXIN1-GFP cells expressing a Tnks shRNA. **c.** AXIN1-GFP stabilization following incubation with compounds for 24h as indicated (n = 3, error bars = SEM). GFP stabilization is quantified as the increase in fluorescence over DMSO and normalized to the highest value. **Boxed:** AXIN1-GFP stabilization of inactive compounds normalized to maximum GFP increase following treatment with XAV.

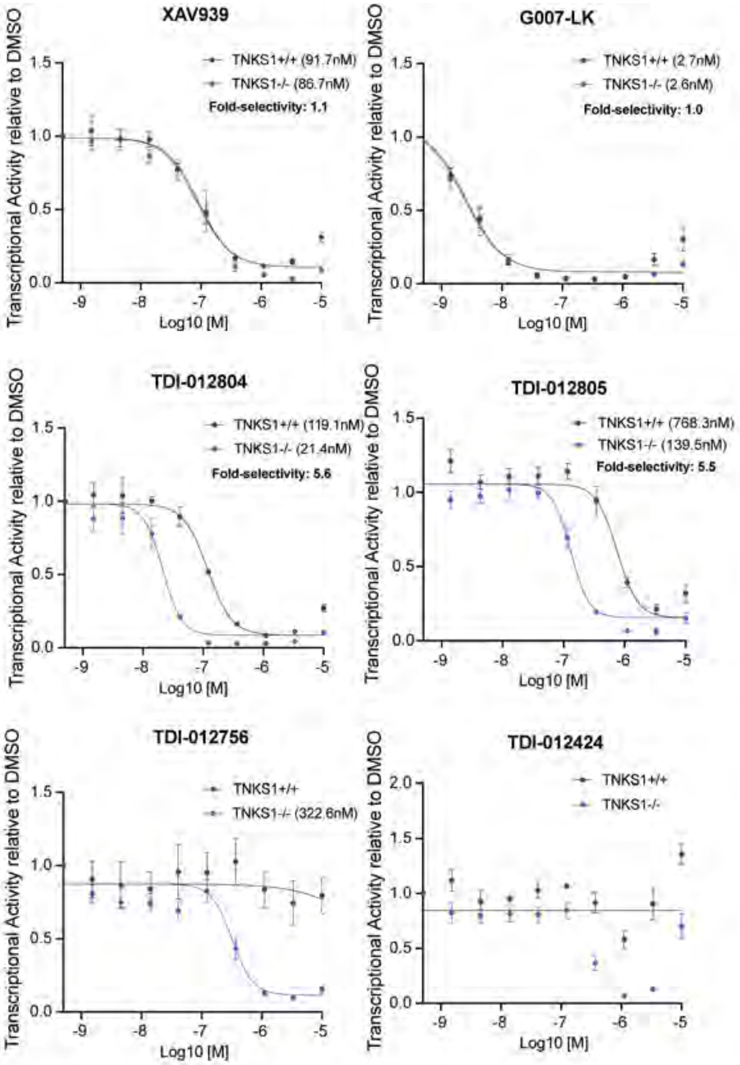


**Figure S8. AXIN1 stabilization and WNT inhibition with hit compounds**

Dose response curves of transcriptional activity measured using a TOPFlash reporter expressed in isogenic DLD1 cell lines (n = 3-4, error bars = SEM). Luciferase is normalized to DMSO.

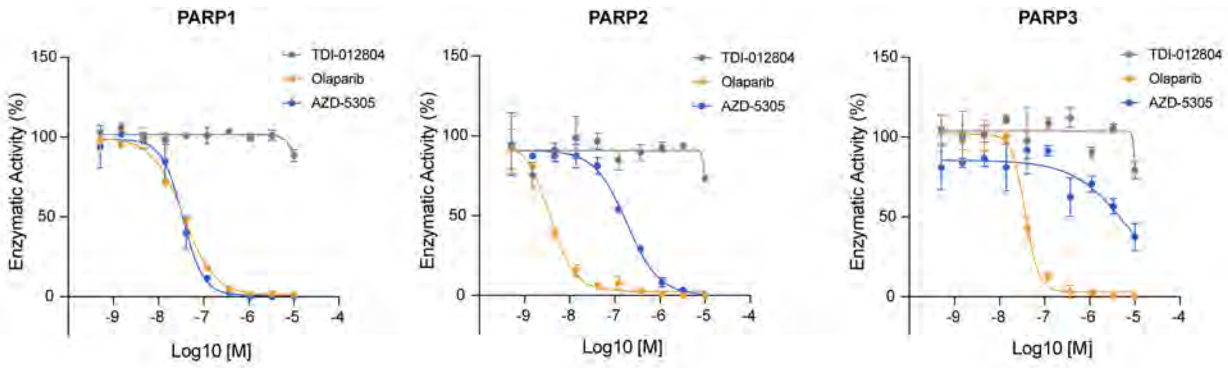


**Figure S9. TDI-012804 does not inhibit PARP1/2/3**
Dose response curves from in vitro PARylation assays measuring the enzymatic activity of PARP1, PARP2, and PARP3 incubated with TDI-012804, Olaparib, or AZD-5305 as indicated (n = 2, error bars = SD). Enzymatic activity is normalized to DMSO.

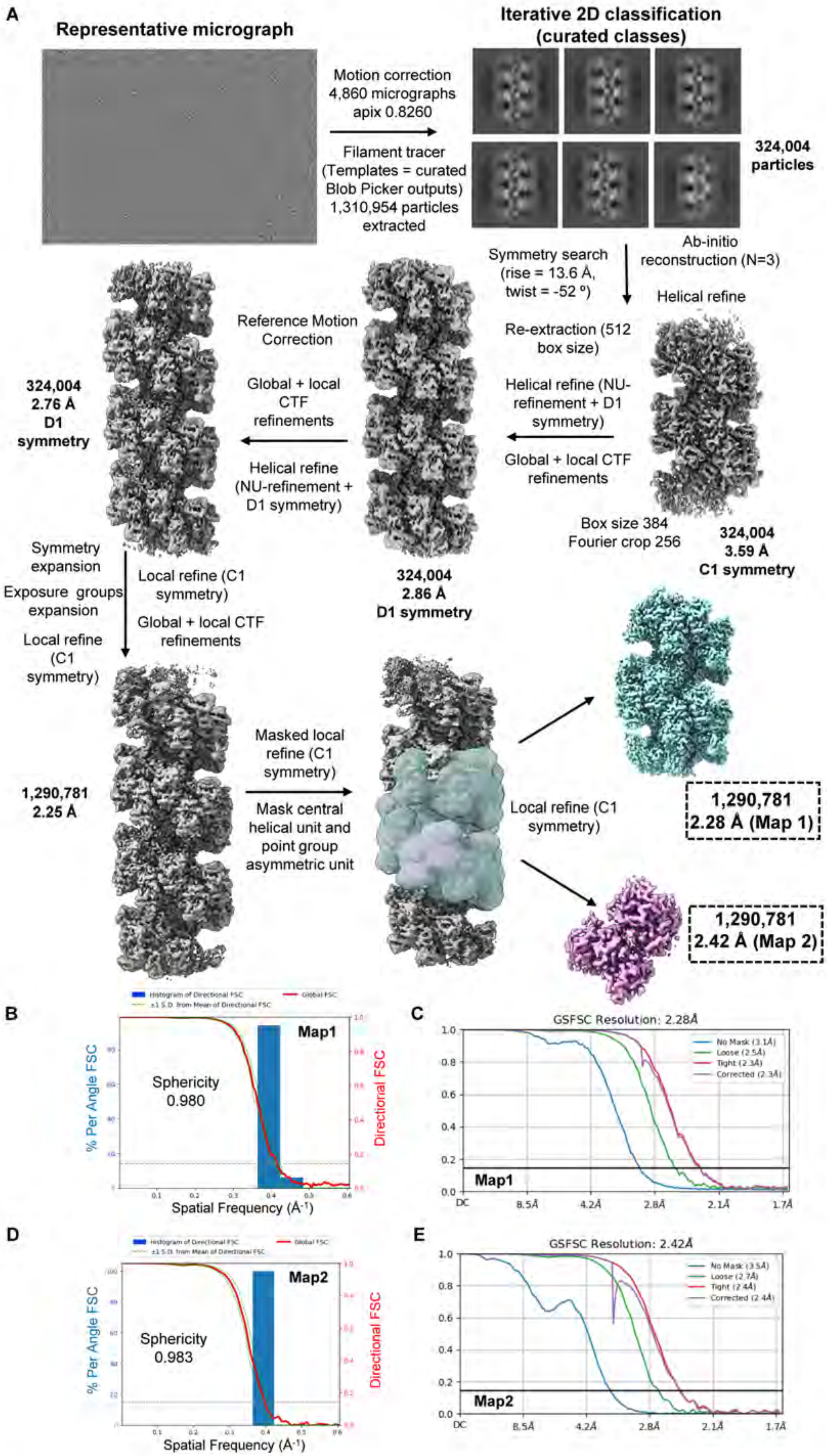


**Figure S10. Cryo-EM processing for the +2804 dataset**

A) Cryo-EM processing pipeline. B) 3D-FSC plot for map1. C) Gold-standard FSC plots for map1. D) 3D-FSC plot for map2. E) Gold-standard FSC plots for map2.

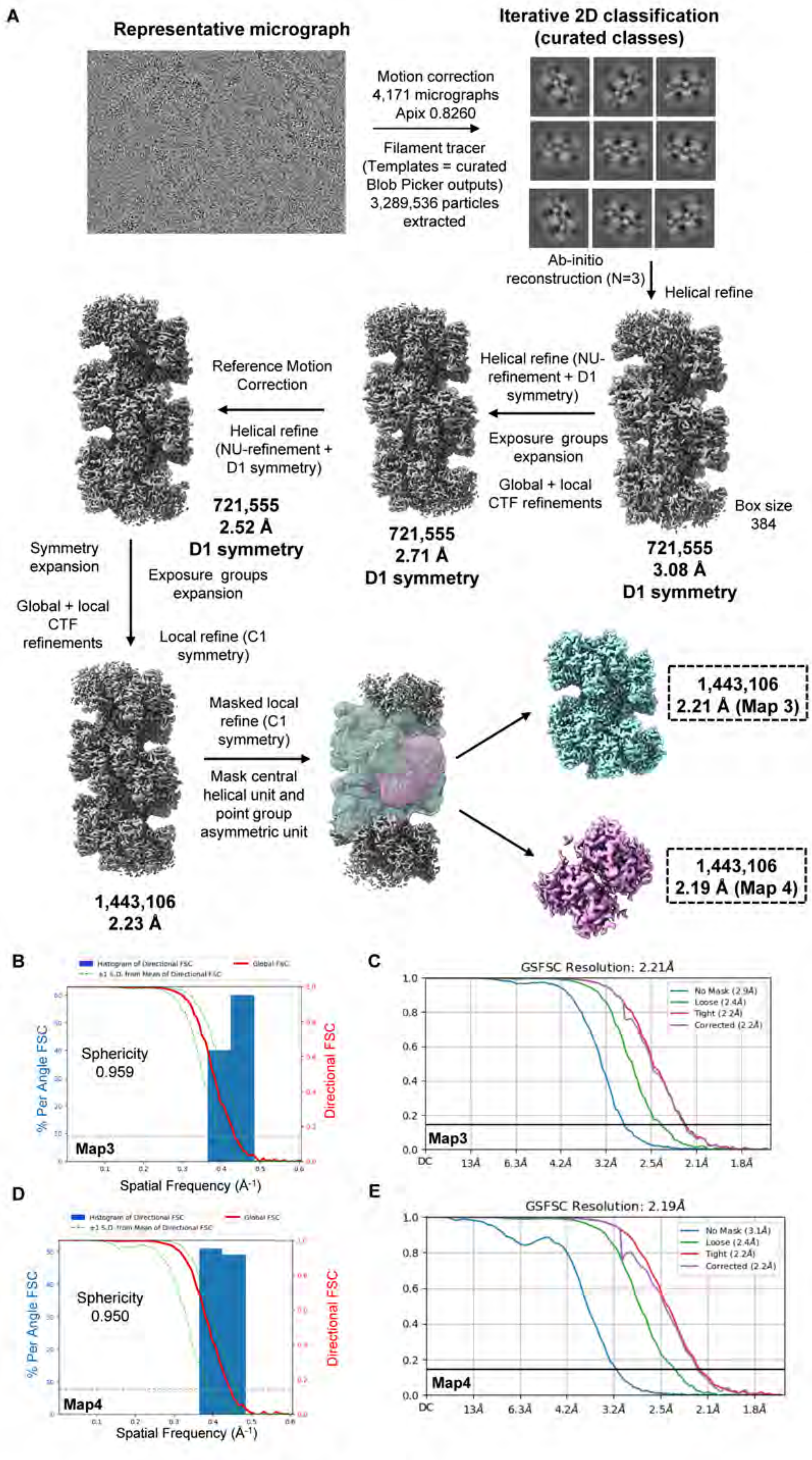


**Figure S11. Cryo-EM processing for the +XAV dataset**

A) Cryo-EM processing pipeline. B) 3D-FSC plot for map3. C) Gold-standard FSC plots for map3. D) 3D-FSC plot for map4. E) Gold-standard FSC plots for map4.

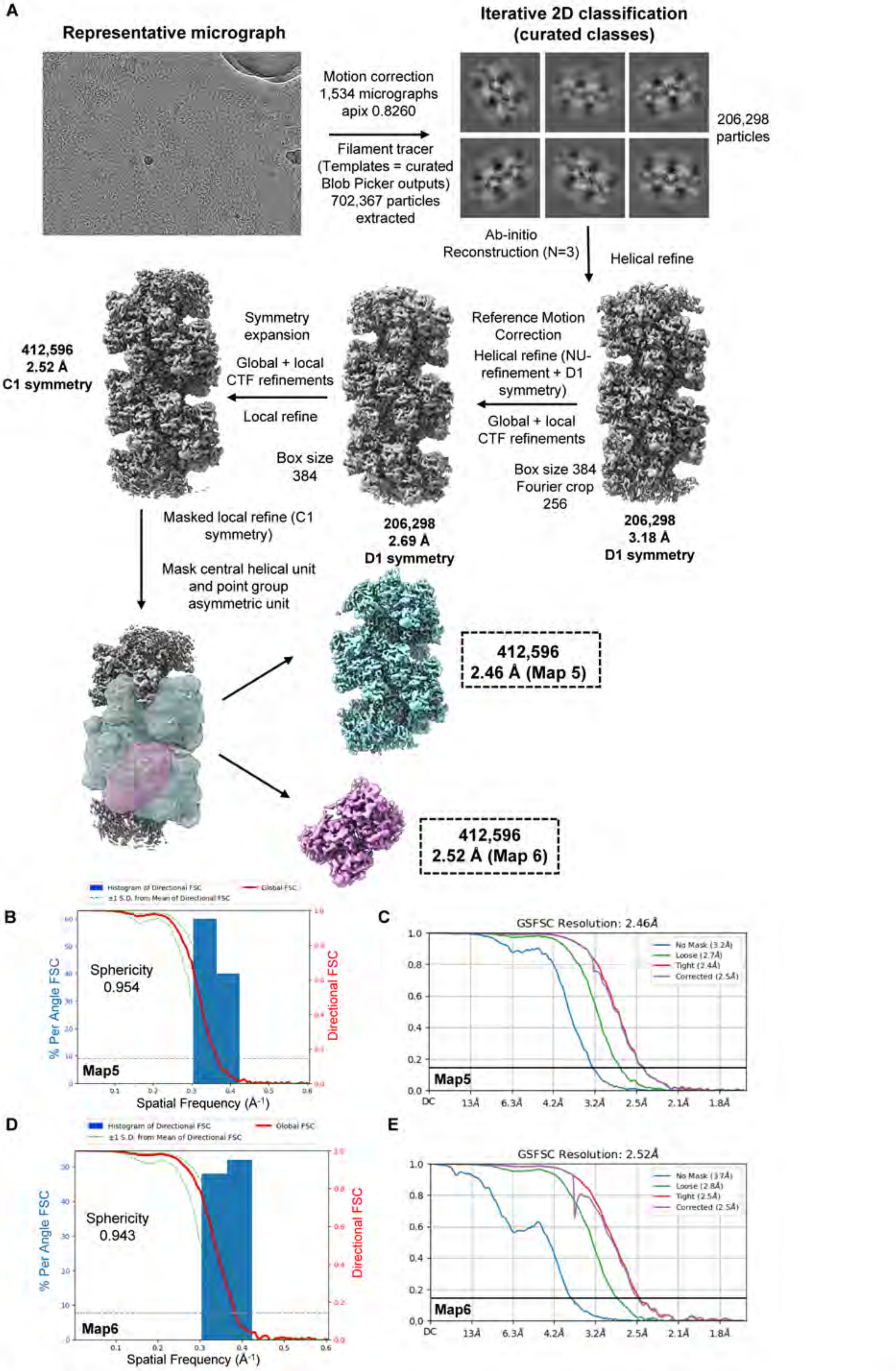


**Figure S12. Cryo-EM processing for the apo dataset**

A) Cryo-EM processing pipeline. B) 3D-FSC plot for map5. C) Gold-standard FSC plots for map5. D) 3D-FSC plot for map6. E) Gold-standard FSC plots for map6.

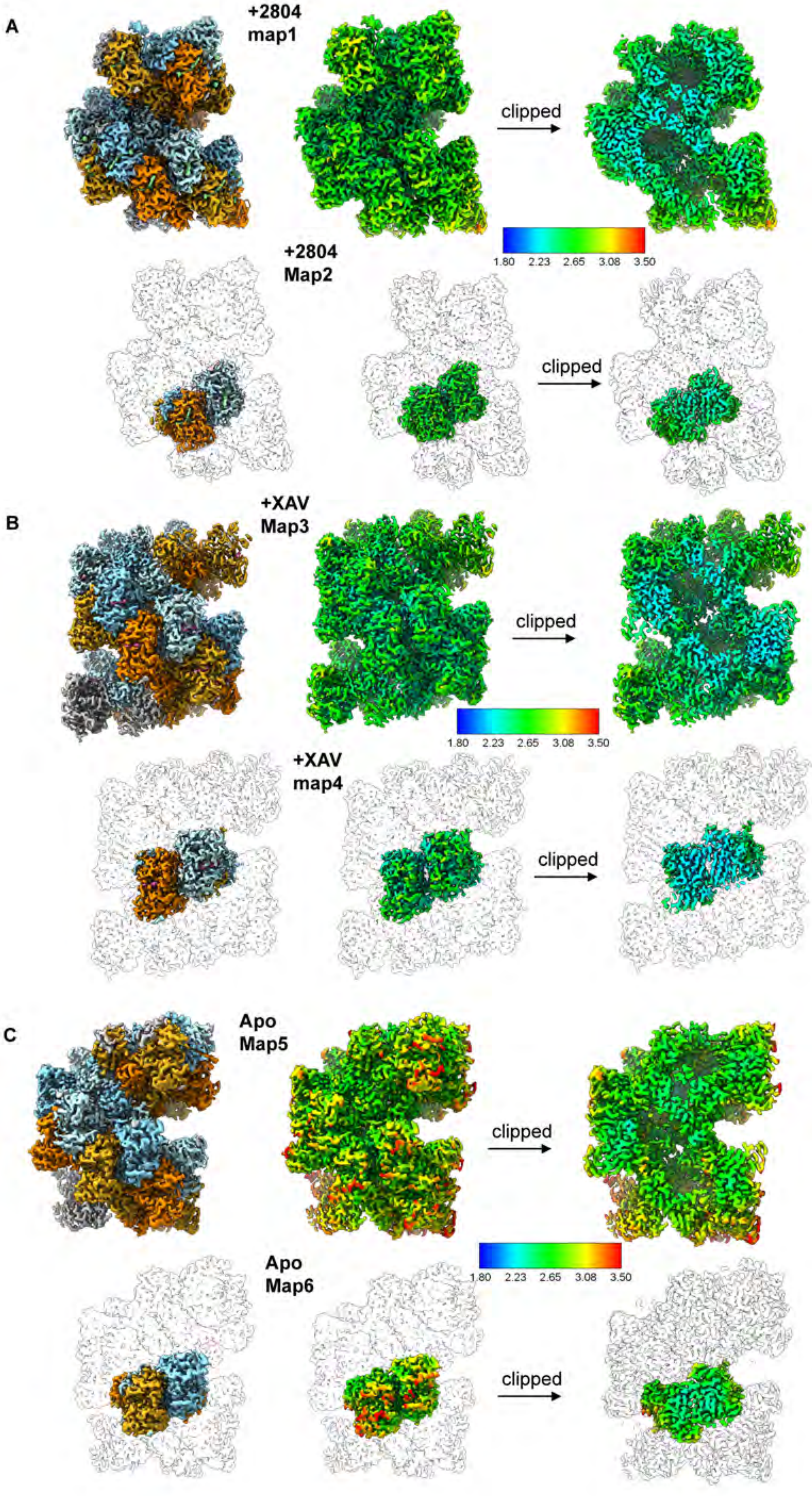


**Figure S13. Local resolution analysis of the final cryo-EM maps.**

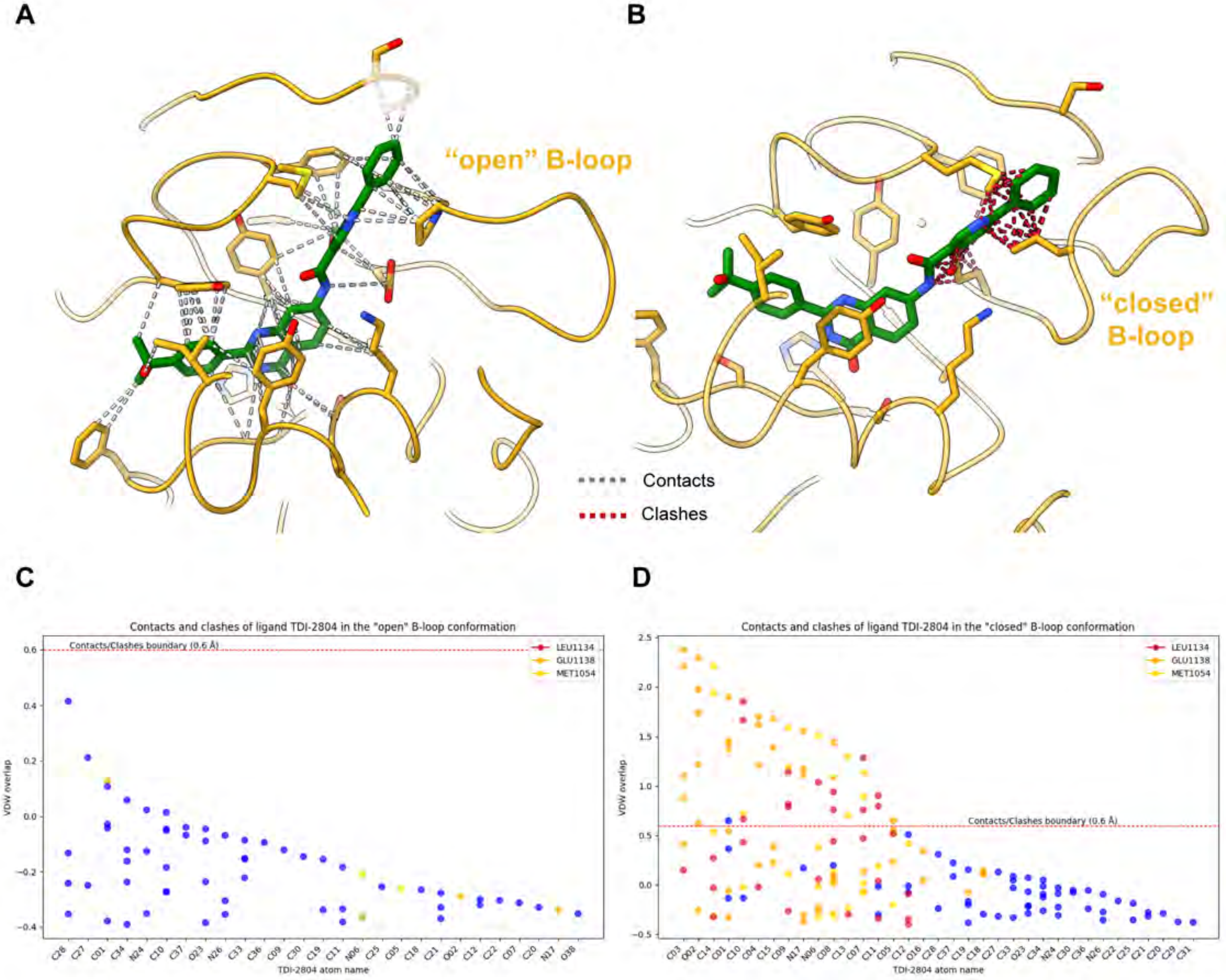


**Figure S14.**

**a.** Analysis of TDI-2804 contacts and clashes with active-site residues (interactions are denoted as dashed gray lines). **b.** Analysis of TDI-2804 clashes with active-site residues when modelled bound to the "closed" B-loop state (clashes are denoted as dashed red lines). **c.** Scatter plot depicting the VDW overlap between the TDI-2804 atoms and surrounding protein residues, plotted interactions from A). **d.** Scatter plot depicting the VDW overlap between the TDI-2804 atoms and surrounding protein residues, plotted interactions from B)

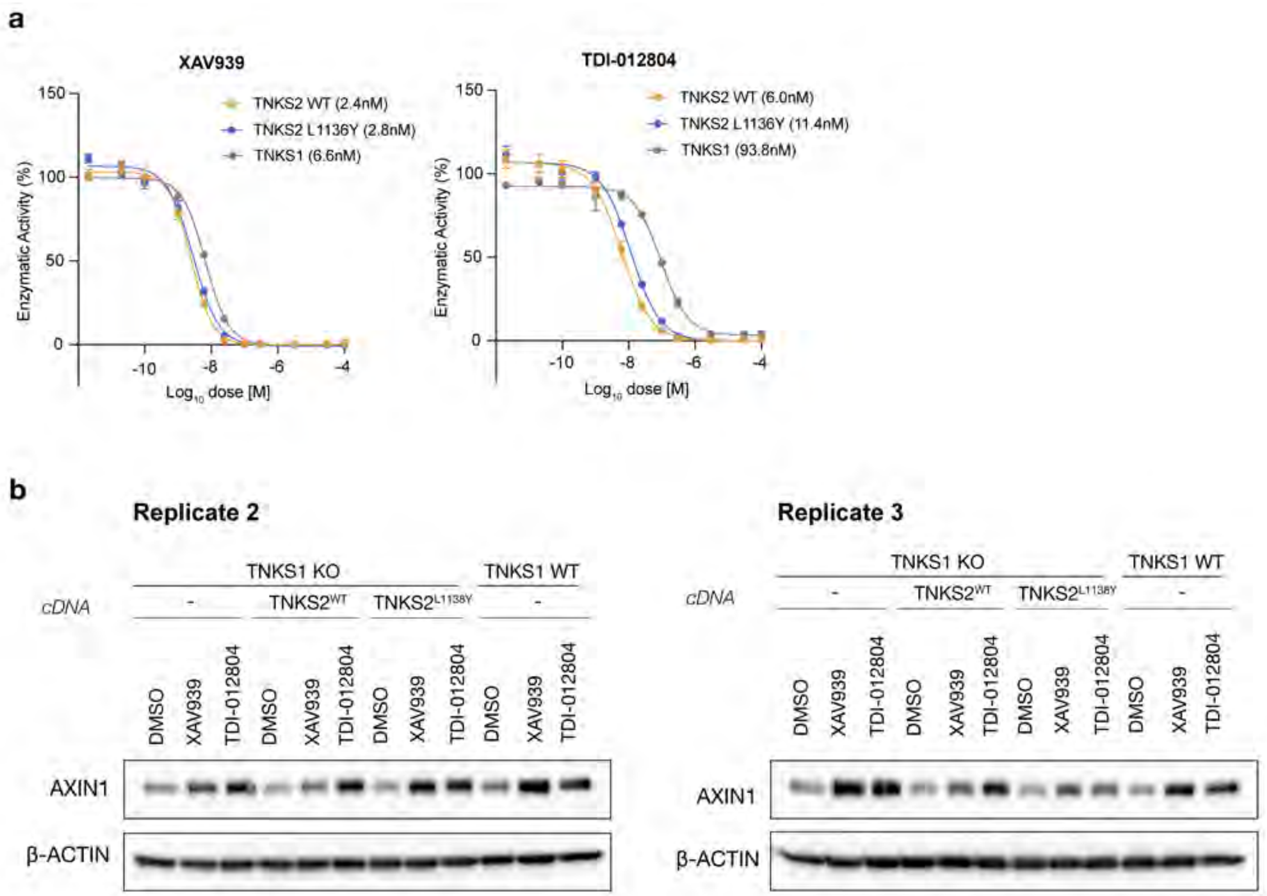


**Figure S15. L1136Y mutation reduces TNKS2 selectivity of TDI-012804**

a. Dose response curves from in vitro PARylation assays measuring the enzymatic activity of TNKS1, $TNKS2^{WT}$, and $TNKS2^{L1136Y}$ incubated with XAV939 or TDI-012804 as indicated (n = 3, error bars = SEM). Enzymatic activity is normalized to DMSO. **b.** Additional replicates of western blot shown in Figure 3G and quantified in Figure 3H.

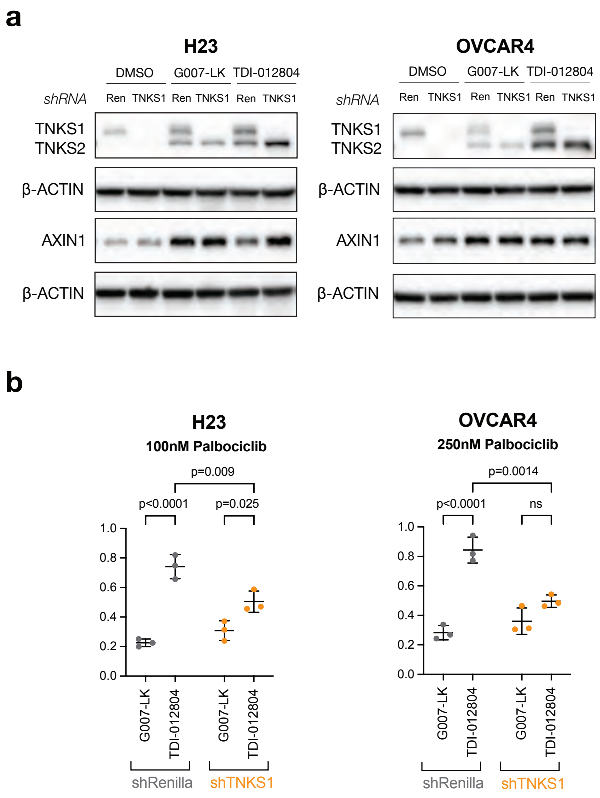


**Figure S16. TDI-012804 selectively reduces proliferation with TNKS1 KD in 8p diploid cell lines.**
**a.** Western blot showing TNKS1/2 and AXIN1 protein levels in H23 and OVCAR3 cells expressing shRenila or shTNKS1 treated with G007-LK (250nM), or TDI-012804 (250nM) for 24h. **b.** Quantification of H23 and OVCAR4 colony forming assays treated with 250nM G007-LK or 250nM TDI-012804 in combination with Palbociclib. (n = 3, mean with SD, two-way ANOVA with Tukey correction).

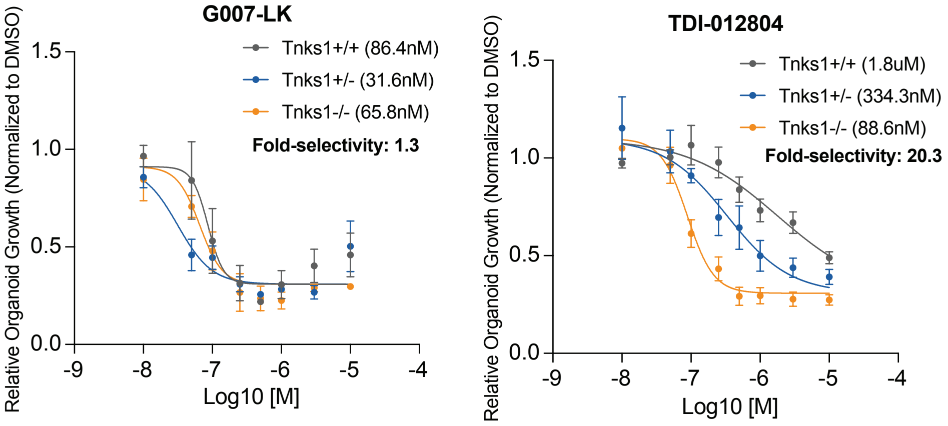


**Figure S17. TDI-012804 selectively reduces cell viability**

Dose response curve showing quantitation of relative organoid area from incucyte imaging of Apc$^{Q1405X}$ small intestinal organoids treated with G007-LK or TDI-012804 for 6 days (n = 3, error bars = SEM). Organoid area is normalized to DMSO.

#### Uncropped Western Blots

##### From Fig 1c

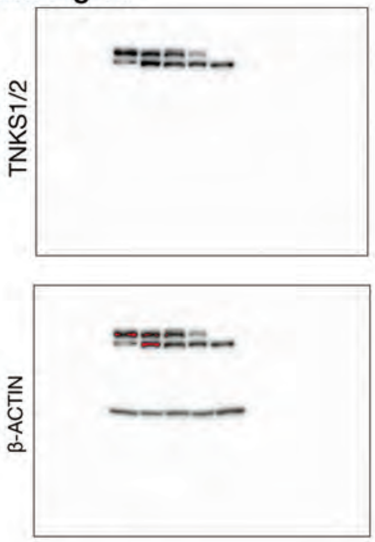


##### From Fig 1d

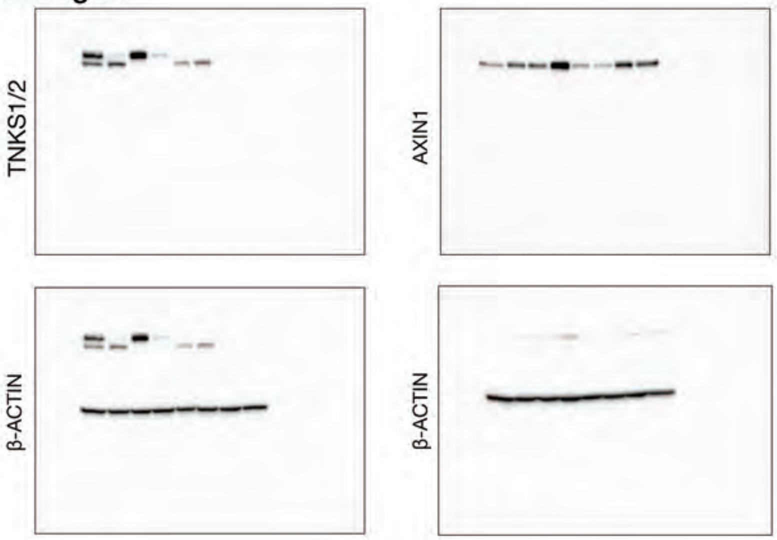


##### From Fig 1e

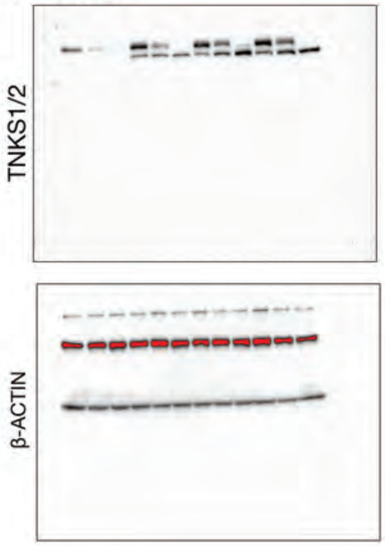


##### From Fig 1f

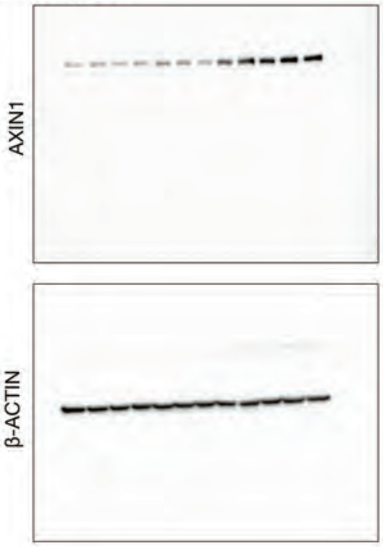


##### From Fig S3a

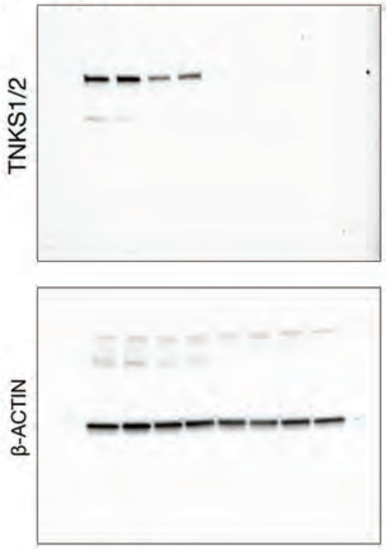


##### From Fig 2f

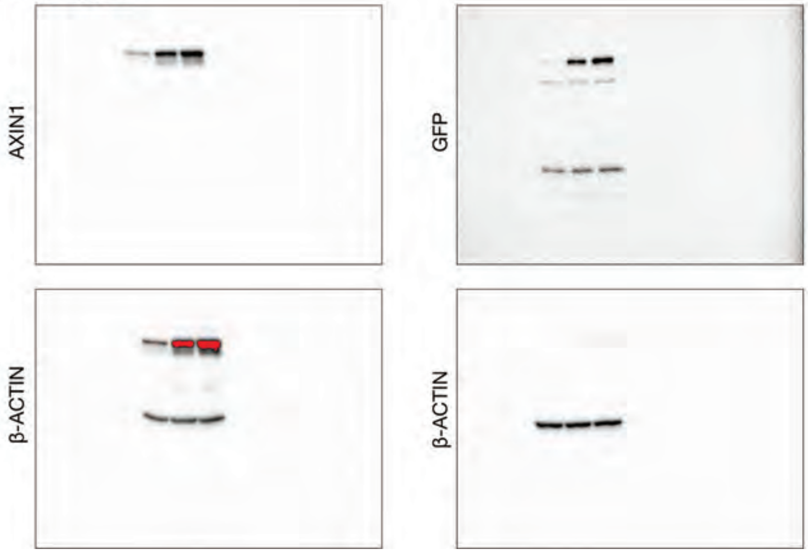


##### From Fig 2j

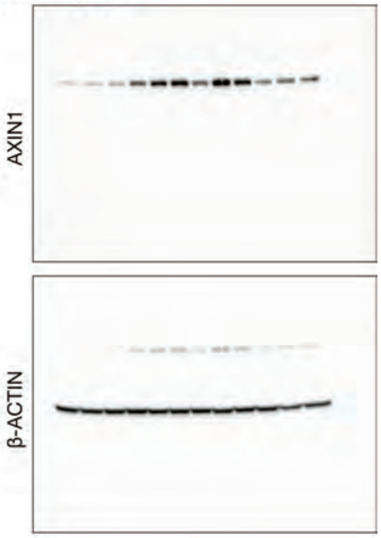

#### Uncropped Western Blots 2

##### From Fig S7b

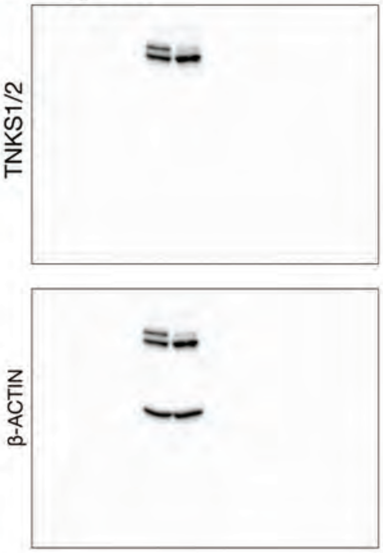


##### From Fig 3g

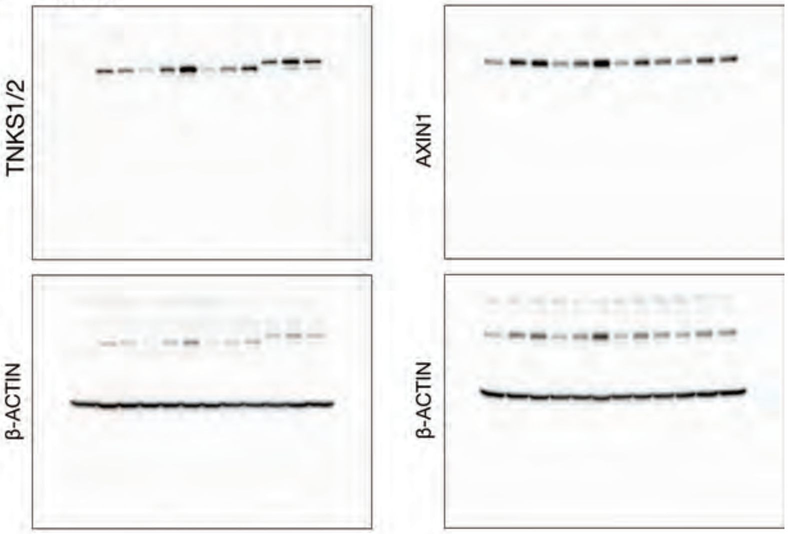


##### From Fig S15b

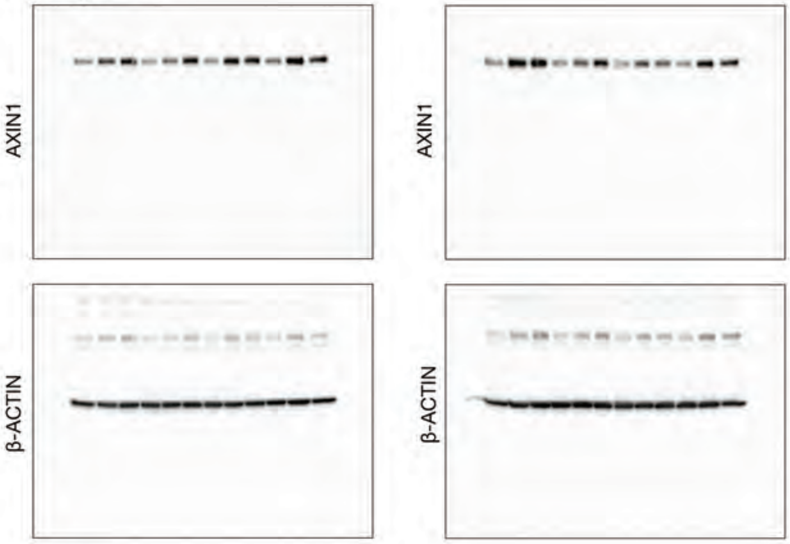


##### From Fig 5a

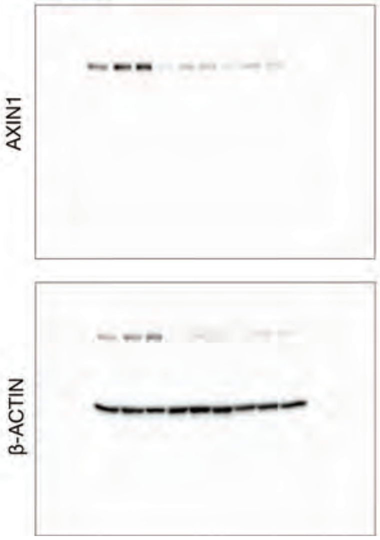

### Uncropped Western Blots 3

#### From Fig S17a

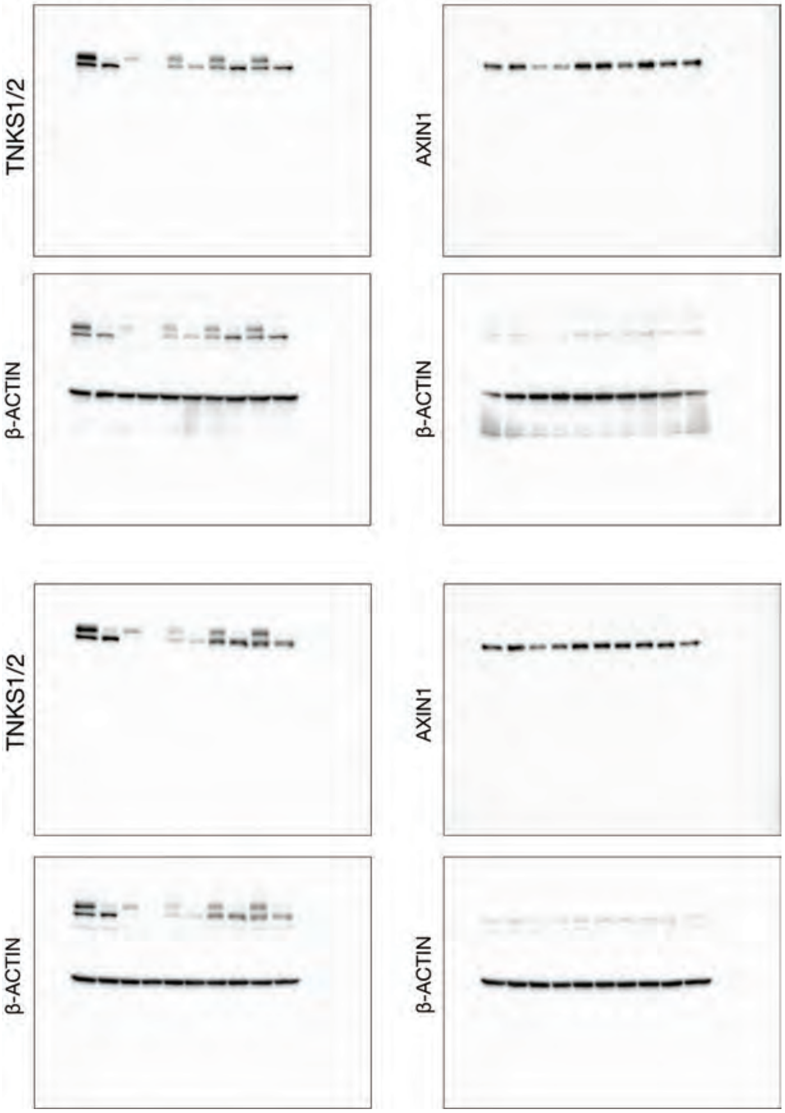
