## Supplementary_Methods for "A potent and selective TNKS2 inhibitor for tumor-selective WNT suppression"

**TDI-012727**

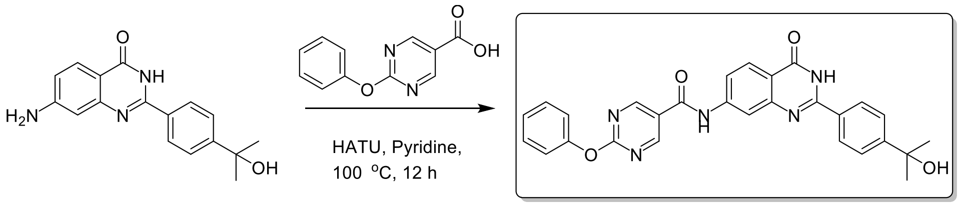


**Step 1** **T051T-200930-7**

#### Step 1 (ET22458-1288):

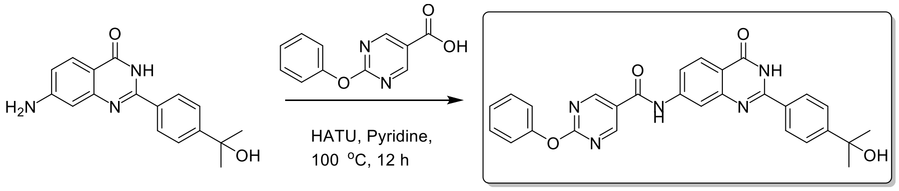


**Step 1** **T051T-200930-7**

##### N-[2-[4-(1-hydroxy-1-methyl-ethyl)phenyl]-4-oxo-3H-quinazolin-7-yl]-2-phenoxy-pyrimidine-5-carboxamide

A mixture of 7-amino-2-[4-(1-hydroxy-1-methyl-ethyl)phenyl]-3H-quinazolin-4-one (80 mg, 270.88 umol, 1 eq), 2-phenoxypyrimidine-5-carboxylic acid (58.56 mg, 270.88 umol, 1 eq) and HATU (103.00 mg, 270.88 umol, 1 eq) in Py. (1 mL) was stirred at 100 °C for 12 h. The reaction mixture was concentrated under reduced pressure to give a residue. The residue was purified by prep-HPLC (column: Phenomenex Gemini-NX 150*30mm*5um;mobile phase:[water(0.05%$NH_3H_2O$+10mM $NH_4HCO_3$)-ACN];B%: 30%-70%, 8 min). Compound N-[2-[4-(1-hydroxy-1-methyl-ethyl)phenyl]-4-oxo-3H-quinazolin-7-yl]-2-phenoxy-pyrimidine-5-carboxamide (16.6 mg, purity 96.92%) was obtained as white solid.

**$^1$H NMR:** ET22458-1288-p1j1 (400MHz, DMSO-$d_6$)

δ 10.83 (s, 1H), 9.16 (s, 2H), 8.27 (br. s, 1H), 8.16-8.11 (m, 3H), 7.81-7.78 (m, 1H), 7.67-7.61 (m, 2H), 7.55-7.45 (t, 3H), 7.33-7.31 (m, 1H), 7.31-7.25 (m, 2H), 5.17 (br. s, 2H), 1.46 (br. s, 6H).

**LCMS:** ET22458-1288-p1j1 (M+H$^+$): 494.2 @ 2.242 min (5-95% ACN in $H_2O$, 6 min)

#### TDI-012465

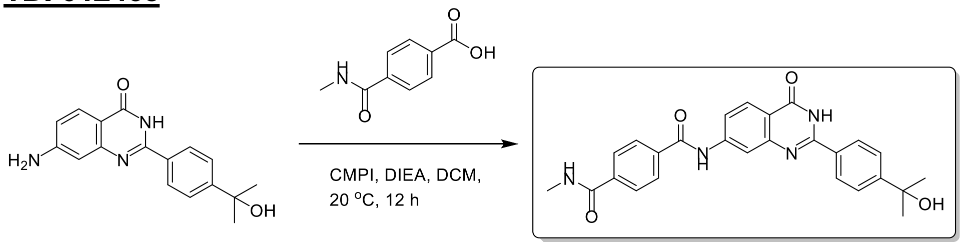


Step 1                T051T-200710-13

#### Step 1 (ET35127-341):

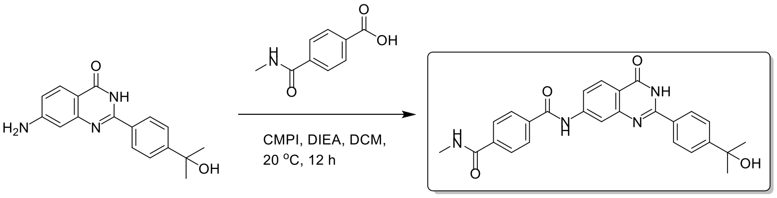


Step 1                T051T-200710-13

**N1-[2-[4-(1-hydroxy-1-methyl-ethyl)phenyl]-4-oxo-3H-quinazolin-7-yl]-N4-methyl-terephthalamide**

A mixture of 7-amino-2-[4-(1-hydroxy-1-methyl-ethyl)phenyl]-3H-quinazolin-4-one (0.1 g, 338.60 umol, 1 *eq*), 4-(methylcarbamoyl)benzoic acid (91.00 mg, 507.90 umol, 1.5 *eq*), 2-chloro-1,3-dimethyl-4,5-dihydroimidazol-1-ium;chloride (114.48 mg, 677.20 umol, 2 *eq*), DIPEA (109.40 mg, 846.50 umol, 147.44 uL, 2.5 *eq*) in DCM (1 mL) was degassed and purged with $N_2$ for 3 times, and then the mixture was stirred at 20 °C for 12 h under $N_2$ atmosphere. The mixture was concentrated in vacuum. The residue was purified by prep-HPLC (neutral condition: column: Waters Xbridge Prep OBD C18 150*40mm*10um; mobile phase: [water(10mM $NH_4HCO_3$)-ACN]; B%: 10%-40%, 8 min). Compound N1-[2-[4-(1-hydroxy-1-methyl-ethyl)phenyl]-4-oxo-3H-quinazolin-7-yl]-N4-methyl-terephthalamide (20.6 mg, 40.48 umol, 11.95% yield, 89.7% purity) was obtained as a yellow oil.

**LCMS:** ET35127-341-P1K2 ($M+H^+$):457.2 @ 1.889 min (5-95 % ACN in $H_2O$, 6 min)

**$^1$H NMR:** ET35127-341-p1g1 (400 MHz, DMSO-d6)

δ 12.41 (br dd, *J* = 1.5, 5.5 Hz, 1H), 10.75 (br s, 1H), 8.70-8.59 (m, 1H), 8.35-8.29 (m, 1H), 8.20-8.11 (m, 3H), 8.09-7.98 (m, 3H), 7.85 (br dd, *J* = 1.6, 8.7 Hz, 1H), 7.71-7.59 (m, 2H), 5.19 (s,1H), 2.87-2.76 (m, 3H), 1.47 (br s, 6H).

#### TDI-012747

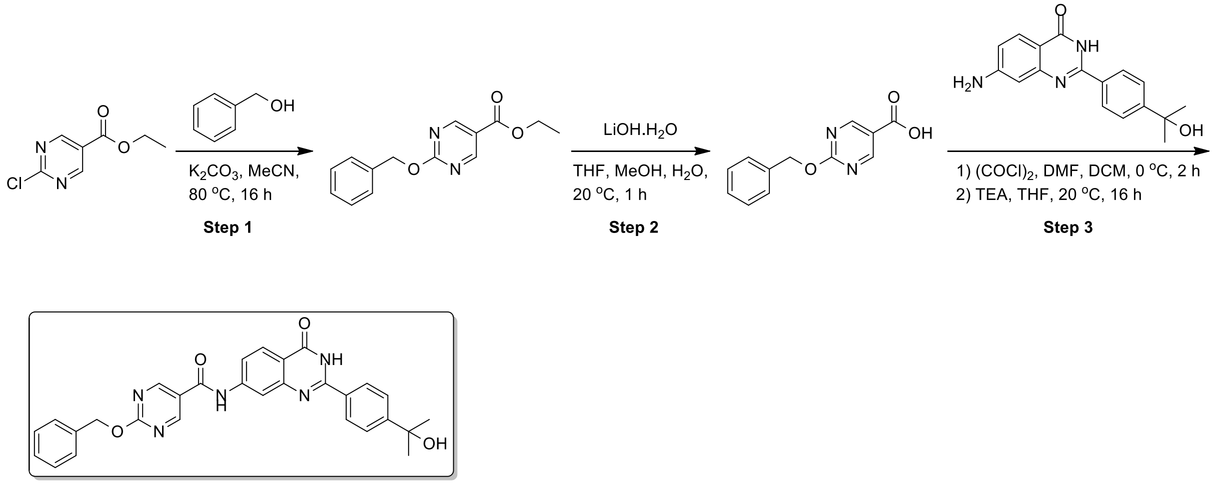


T051T-200930-8

#### Step 1 (ET27680-409):

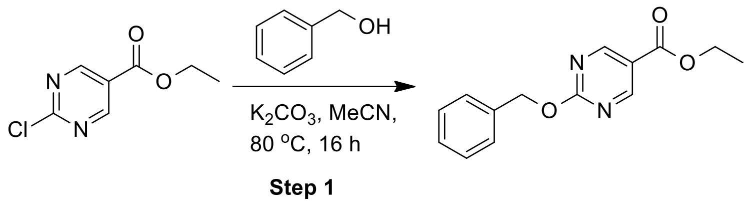


**ethyl 2-benzyloxypyrimidine-5-carboxylate**
A mixture of phenylmethanol (695.44 mg, 6.43 mmol, 668.69 uL, 1.2 eq), ethyl 2-chloropyrimidine-5-carboxylate (1 g, 5.36 mmol, 1 eq), $K_2CO_3$ (1.11 g, 8.04 mmol, 1.5 eq) in MeCN (10 mL) was degassed and purged with $N_2$ for 3 times, and then the mixture was stirred at 80 °C for 16 h under $N_2$ atmosphere. The mixture was poured into $H_2O$ (50 mL), extrated with EtOAc (20 mL*3). The combined organic layer was washed with brine (50 mL), dried over $Na_2SO_4$, filtered and concentrated under reduced pressure. The residue was purified by column chromatography ($SiO_2$, Petroleum ether/Ethyl acetate = 1/0 to 10/1) to give ethyl 2-benzyloxypyrimidine-5-carboxylate (0.43 g, crude) as a white solid.

**LCMS:** ET27680-409-P1A $(M+H^+)$:259.1@ 0.762 min (5_95AB_2min-220-254, 1.5 min)

#### Step 2 (ET27680-414):

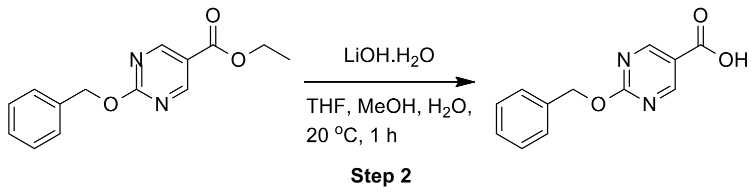

**2-benzyloxypyrimidine-5-carboxylic acid**

To a solution of ethyl 2-benzyloxypyrimidine-5-carboxylate (230 mg, 890.53 umol, 1 eq) in THF (3 mL), MeOH (1 mL) and $H_2O$ (1 mL) was added LiOH.$H_2O$ (112.10 mg, 2.67 mmol, 3 eq). The mixture was stirred at 20 °C for 1 h. The mixture was concentrated under reduced pressure, and then the residue was adjust pH=2 with HCl (2 N). The mixture was filtered and the filtere cake was concentrated under reduced pressure to give methyl 2-benzyloxypyrimidine-5-carboxylic acid (0.11 g, crude) as a white solid.

**LCMS:** ET27680-414-P1A (M+$H^+$):231.1@ 0.628 min (5_95AB_2min-220-254, 1.5 min)

**Step 3 (ET27680-435&436):**

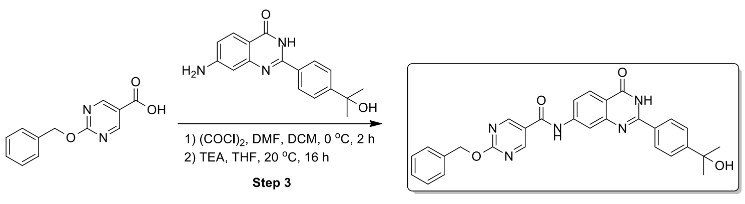

T051T-200930-8

**2-benzyloxy-N-[2-[4-(1-hydroxy-1-methyl-ethyl)phenyl]-4-oxo-3H-quinazolin-7-yl]pyrimidine-5-carboxamide**

To a solution of 2-benzyloxypyrimidine-5-carboxylic acid (0.1 g, 434.37 umol, 1 eq) in DCM (2 mL) was added $(COCl)_2$ (82.70 mg, 651.55 umol, 57.03 uL, 1.5 eq) and DMF (3.17 mg, 43.44 umol, 3.34 uL, 0.1 eq). The mixture was stirred at 0 °C for 2 h. The mixture was concentrated under reduced pressure. Compound 2-benzyloxypyrimidine-5-carbonyl chloride (0.1 g, crude) was obtained as a white solid. The crude product was used into the next step without further purification. A mixture of 2-benzyloxypyrimidine-5-carbonyl chloride (0.1 g, 402.15 umol, 1 eq), 7-amino-2-[4-(1-hydroxy-1-methyl-ethyl)phenyl]-3H- quinazolin-4-one (142.52 mg, 482.58 umol, 1.2 eq), $Et_3N$ (40.69 mg, 402.15 umol, 55.97 uL, 1 eq) in THF (4 mL) was degassed and purged with $N_2$ for 3 times, and then the mixture was stirred at 20 °C for 16 h under $N_2$ atmosphere. The mixture was concentrated under reduced pressure. The residue was purified by prep-HPLC (neutral condition; column: Phenomenex Gemini-NX 80 * 40 mm* 3 um; mobile phase: [water (10mM $NH_4HCO_3$) - ACN]; B%: 30% -50%, 8 min) to give 2-benzyloxy-N-[2-[4-(1-hydroxy-1-methyl-ethyl)phenyl]-4-oxo-3H- quinazolin-7-yl]pyrimidine-5-carboxamide (8 mg, 15.51 umol, 3.86% yield, 98.39% purity) as a white solid.

**LCMS:** ET27680-436-P1C $(M+H^+)$:508.2@ 2.353 min (5_95AB_6min-220, 4.5 min)

**$^1$H NMR:** ET27680-436-P1A (400 MHz, DMSO)

δ 12.32 (br s, 1H), 10.77 (s, 1H), 9.17 (s, 2H), 8.26 (s, 1H), 8.14 (d, *J* = 8.5 Hz, 3H), 7.84-7.74 (m, 1H), 7.63 (d, *J* = 8.4 Hz, 2H), 7.50 (br d, *J* = 7.0 Hz, 2H), 7.45-7.32 (m, 3H), 5.60-5.44 (m, 2H), 5.18 (s, 1H), 1.47 (s, 6H).

**TDI-012756**

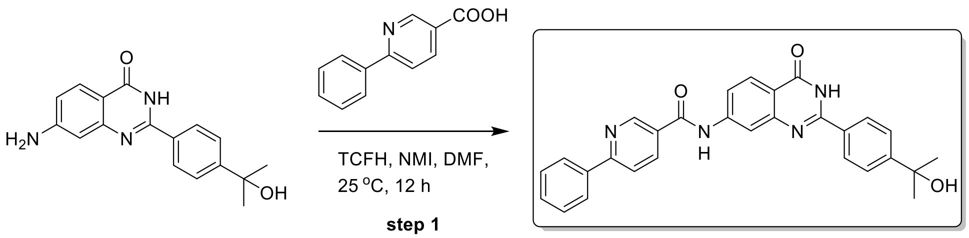


**T051T-200930-2**

#### Step 1 (ET22458-1289):

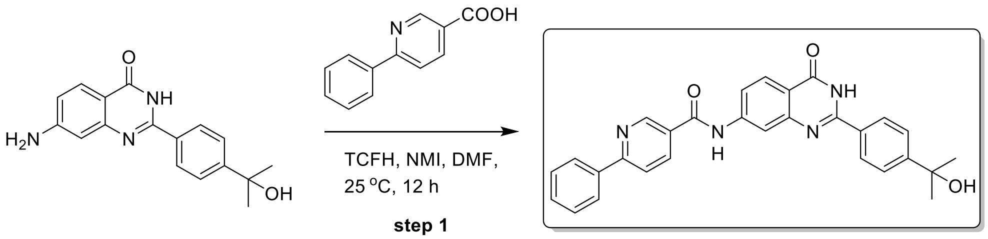


**T051T-200930-2**

##### N-[2-[4-(1-hydroxy-1-methyl-ethyl)phenyl]-4-oxo-3H-quinazolin-7-yl]-6-phenyl-pyridine-3-carboxamide

A mixture of 7-amino-2-[4-(1-hydroxy-1-methyl-ethyl)phenyl]-3H-quinazolin-4-one (0.1 g, 338.60 umol, 1 eq), 6-phenylpyridine-3-carboxylic acid (80.94 mg, 406.32 umol, 1.2 eq) and 6-phenylpyridine-3-carboxylic acid (80.94 mg, 406.32 umol, 1.2 eq) in DMF (2 mL) was added [chloro(dimethylamino)methylene]-dimethyl-ammonium;hexafluorophosphate (104.50 mg, 372.46 umol, 1.1 eq) and 1-methylimidazole (69.50 mg, 846.50 umol, 67.48 uL, 2.5 eq). The mixture was stirred at 25 °C for 12 h. The mixture was filtered and concentrated in vacuum. The mixture was purified by prep-HPLC (column: Phenomenex Gemini-NX C18 75*30mm*3um; mobile phase: [water (0.05% $NH_3H_2O$ + 10mM $NH_4HCO_3$)-ACN]; B%: 30%-60%, 8min). Compound N-[2-[4-(1-hydroxy-1-methyl-ethyl)phenyl]-4-oxo-3H-quinazolin-7-yl]-6-phenyl-pyridine-3-carboxamide (28.8 mg, 60.06 umol, 17.74% yield, 99.37% purity) was obtained as white solid.

**$^1$H NMR:** ET22458-1289-p1j3 (400MHz, DMSO-*d6*)
δ 12.38 (br. s, 1H), 10.83 (s, 1H), 9.25 (s, 1H),8.47-8.52 (m, 1H), 8.23 (s, 1H), 8.22-8.16 (m, 3H), 8.16-8.13 (m, 3H), 7.84-7.80 (m, 1H) 7.66-7.64 (m, 2H), 7.63-6.50 (m, 3H), 5.17 (s, 1H), 1.48 (s, 6H).

**LCMS:** ET22458-1289-p1j1 (M+H$^+$): 477.2 @ 2.296 min (5-95% ACN in $H_2O$, 4.5 min)

#### TDI-012424

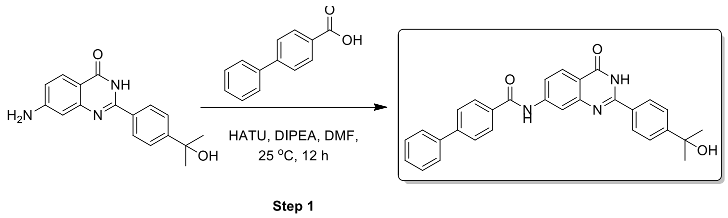


T051T-200710-16

#### Step 1 (ET35626-173)

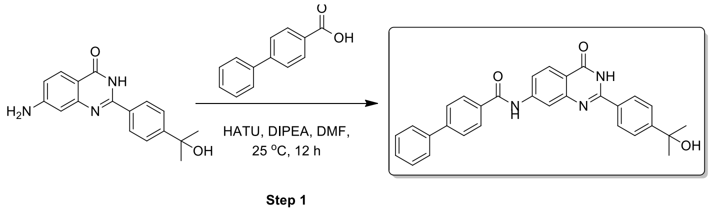


T051T-200710-16

**N-[2-[4-(1-hydroxy-1-methyl-ethyl)phenyl]-4-oxo-3H-quinazolin-7-yl]-4-phenyl-benzamide**

To a solution of 7-amino-2-[4-(1-hydroxy-1-methyl-ethyl)phenyl]-3H-quinazolin-4-one (50 mg, 169.30 umol, 1 eq), 4-phenylbenzoic acid (50.34 mg, 253.95 umol, 1.5 *eq*), HATU (96.56 mg, 253.95 umol, 1.5 *eq*) in DMF (1 mL) was added DIPEA (54.70 mg, 423.25 umol, 73.72 uL, 2.5 *eq*). The mixture was stirred at 25 °C for 12 hr. The mixture was collected by filtration and the filtrate was concentrated under reduced pressure to give a residue. The residue was purified by prep-HPLC (HCl condition, column: Phenomenex Luna C18 75*30mm*3um; mobile phase: [water (0.2%FA)-ACN]; B%: 40%-70%,8 min). Compound N-[2-[4-(1-hydroxy-1-methyl-ethyl)phenyl]-4-oxo-3H-quinazolin-7-yl]-4-phenyl-benzamide (11.6 mg, 21.76 umol, 12.85% yield, 89.21% purity) was obtained as a white solid.

**LCMS:** ET35626-173-P1C ($M+H^+$):476.1 @ 2.611 min (5-95 % ACN in $H_2O$, 6 min)

**$^1$H NMR:** ET35626-173-P1A (400 MHz, DMSO-d6)

δ 10.68 (s, 1H), 8.34 (s, 1H), 8.16-8.09 (m, 5H), 7.88 (br d, *J* = 8.38 Hz, 3H), 7.78 (br d, *J* = 7.25 Hz, 2H), 7.63 (d, *J* = 8.38 Hz, 2H), 7.55-7.49 (m, 2H), 7.46-7.41 (m, 1H), 1.47 (s, 6H).

#### TDI-012804

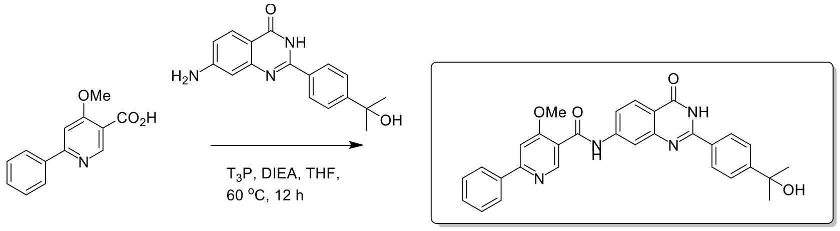


Step 1 T051T-220302-1

#### Step 1 (ET16082-3178):

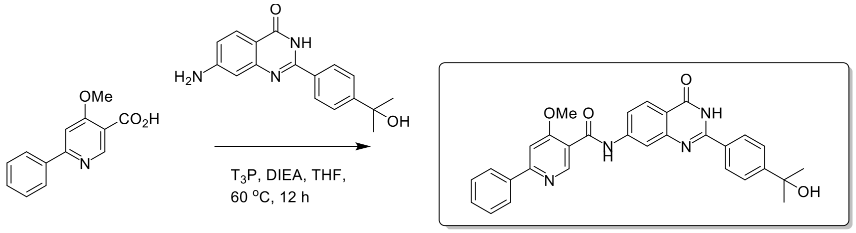


Step 1 T051T-220302-1

**N-[2-[4-(1-hydroxy-1-methyl-ethyl)phenyl]-4-oxo-3H-quinazolin-7-yl]-4-methoxy-6-phenyl-pyridine-3-carboxamide**

A mixture of 7-amino-2-[4-(1-hydroxy-1-methyl-ethyl)phenyl]-3H-quinazolin-4-one (0.5 g, 1.69 mmol, 1 *eq*), 4-methoxy-6-phenyl-pyridine-3-carboxylic acid (388.09 mg, 1.69 mmol, 1 *eq*), $T_3P$ (4.31 g, 6.77 mmol, 4.03 mL, 50% purity, 4 *eq*), DIPEA (1.09 g, 8.46 mmol, 1.47 mL, 5 *eq*) in THF (10 mL) was degassed and purged with $N_2$ for 3 times, and then the mixture was stirred at 60 °C for 12 h under $N_2$ atmosphere. The residue was poured into $H_2O$ (100 mL). The aqueous phase was extracted with ethyl acetate (50 mL * 3). The combined organic phase was washed with brine (60 mL), dried with anhydrous $Na_2SO_4$, filtered and concentrated in vacuum. The residue was purified by prep-HPLC (neutral condition, column: Phenomenex C18 75*30mm*3um; mobile phase: [water ($NH_4HCO_3$) -ACN]; B%: 30%-60%, 8 min). Compound N-[2-[4-(1-hydroxy-1-methyl-ethyl)phenyl]-4-oxo-3H-quinazolin-7-yl]-4-methoxy-6-phenyl-pyridine-3-carboxamide (110 mg, 100% purity) was obtained as a white solid.

**LCMS:** ET16082-3178-P1B ($M+H^+$):507.2 @ 2.591 min (5-95 % ACN in $H_2O$, 6 min)

**$^1$H NMR:** ET16082-3178-P1B (400 MHz, DMSO-d6)

δ 10.62 (s, 1H), 8.76 (s, 1H), 8.25 (s, 1H), 8.21 (br d, 2H), 8.16-8.09 (m, 3H), 7.80-7.71 (m, 2H), 7.63 (d, 2H), 7.57-7.49 (m, 3H), 5.17 (s, 1H), 4.14-4.09 (m, 3H), 1.47 (s, 6H).

### HRMS:

#### Elemental Composition Report

Page 1

##### Single Mass Analysis

Tolerance = 5.0 PPM / DBE: min = -1.5, max = 50.0
Element prediction: Off
Number of isotope peaks used for i-FIT = 3

Monoisotopic Mass, Even Electron Ions
18 formula(e) evaluated with 1 results within limits (up to 50 closest results for each mass)
Elements Used:
C: 0-30 H: 0-26 N: 0-5 O: 0-4

Rui Liang
TDI
C30H26N4O4
TDI_012804 41 (0.934)

NMR Analytical Core Facility
LCT Premier XE

30-Apr-2024
12:23:19

1: TOF MS ES-
5.63e+003

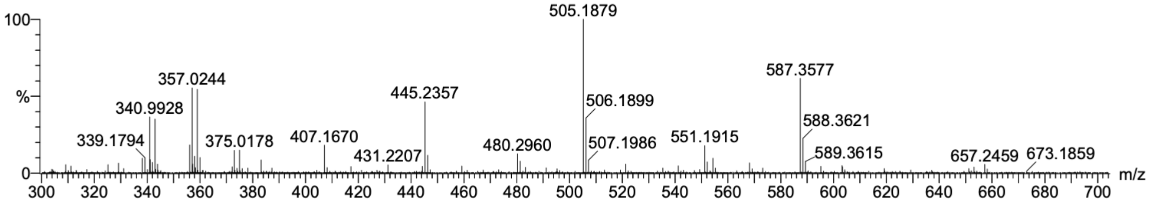


| Mass | Calc. Mass | mDa | PPM | DBE | i-FIT | i-FIT (Norm) | Formula |
|---|---|---|---|---|---|---|---|
| Minimum: | | | | -1.5 | | | |
| Maximum: | | 5.0 | 5.0 | 50.0 | | | |
| 505.1879 | 505.1876 | 0.3 | 0.6 | 20.5 | 122.0 | 0.0 | C30 H25 N4 O4 |

### 13C NMR:

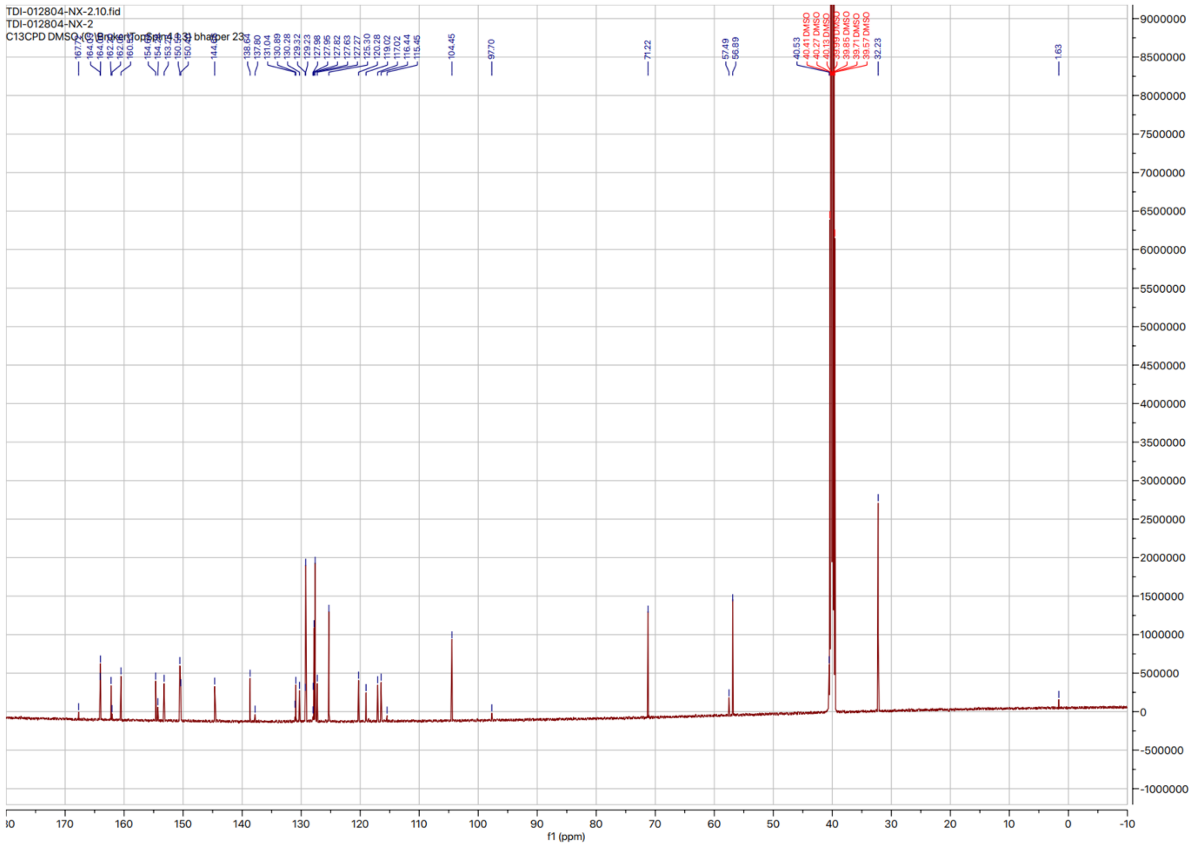
